# Dopaminergic mechanisms of dynamical social specialization in mouse microsocieties

**DOI:** 10.1101/2025.02.10.637303

**Authors:** C. Solié, A. Nicolson, R. Justo, Y. Layadi, B. Morin, C. Batifol, LM Reynolds, T. Le Borgne, SL Fayad, A. Gulmez, J. Allegret--Vautrot, F. de Chaumont, C. Viollet, S. Didienne, N. Debray, J-P. Hardelin, B. Girard, A. Mourot, J. Naudé, F. Marti, B. Delord, P. Faure

**Affiliations:** Brain Plasticity Laboratory, CNRS UMR 8249, ESPCI Paris, PSL Research University, Paris, France; Institut des Systèmes Intelligents et de Robotique (ISIR), Sorbonne Université, CNRS, Paris, France; Montpellier Inserm, CNRS, Université de Montpellier; Institut de Génomique Fonctionnelle, Montpellier, France; Human Genetics and Cognitive Functions, CNRS UMR UMR3571, Pasteur Institute, Paris, France

## Abstract

Social organization and division of labor are fundamental to animal societies, but how do these structures emerge from individual interactions, and what role does neuromodulation play in shaping them? Using behavioral tracking in a semi-natural environment, neural recordings, and computational models integrating reinforcement and social learning, we show that groups of three isogenic mice spontaneously develop specialized roles while solving a foraging task requiring individual decisions under social constraints. Moreover, these roles are shaped by dopaminergic activity in the ventral tegmental area. Strikingly, despite minor sex-differences in behavior when mice were tested alone, male triads formed stable worker-scrounger relationships driven by competition, whereas female triads adopted uniform, cooperative strategies. Model analysis revealed how intra- and inter-sex parameter differences in resource exploitation, combined with contingent and dynamic social interactions, drive behavioral specialization and labor division. Most notably, it highlighted how contingency, amplified by competition, magnifies individual differences and shapes social profiles. The plastic, adaptive nature of social organization within triads was confirmed by manipulating dopaminergic cell activity, which reshaped social roles and altered group structure. Our findings support a feedback loop where social context shapes neural states, which in turn reinforce behavioral specialization and stabilize social structures.

---

Social animals interact and coordinate their behaviors to form social organizations: structured patterns of relationships and interactions that stabilize into collective social forms (e.g., division of labor, norms, institutions) and specialized individual roles^1–5^. A key challenge in behavioral science is deciphering how individual differences in behavior emerge, how they contribute to collective actions, and how they are linked to neural activity. While the broad effects of social organization on individual behavior and specialization are well documented ^6–12^, the underlying cognitive and neurophysiological mechanisms remain largely unexplored. In particular, it is unclear how individual traits shape collective behaviors and social structures. This gap persists largely due to the difficulty of studying such mechanisms in controlled, artificial social settings that also allow for neurobiological investigation.

In social groups, the production of and access to shared resources drives strategies such as competition or cooperation^7,13,14^. This dynamic is exemplified by the producer-scrounger game, where individuals either produce resources (e.g., press a lever to obtain food) or scrounge from others who have already produced them^13,15–19^. Foraging strategy thus refers to the behavioral patterns and decision-making processes individuals adopt to obtain food, balancing effort, risk, and reward^14^. These strategies can range from independent food acquisition to social exploitation of others’ efforts, shaped by ecological constraints and social dynamics. Evolutionary game theory^20^ predicts stable outcomes under such interactions, such as the equilibrium between producer and scrounger strategies, depending on resource availability. However, this theory typically assumes behaviors are predetermined and stable^21^, overlooking the internal mechanisms that guide an individual’s behavior, including the neural and cognitive processes which grant adaptation and learning-based flexible strategies^21–23^. Here, we hypothesize that social foraging strategies, including producer-scrounger dynamics, emerge flexibly through socially-constrained reinforcement learning, with dopaminergic systems playing a central role in behavioral specialization. As social organization forms and norms develop, individuals are expected to adjust their actions based on expected payoffs, which affect both their neural activity and activity-dependent learning processes. In turn, these learning-induced changes in neural activity and in cognitive decisions would shape how individuals behave in social context^24^.

Advances in tracking mice in semi-natural environments^25–30^ provide a unique opportunity to explore how social organization emerges from learning. By continuously monitoring individual behavior and interactions, we investigated how foraging specialization develops in small groups (n=3) housed in a controlled semi-natural setting. Males rapidly adopted distinct roles with divisions of labor, while females maintained uniform strategies. Modeling showed that learning and cognitive processes drive specialization, with exploration-exploitation trade-offs shaping social roles. Dopaminergic manipulations confirmed dopamine’s key role in structuring social organization and reinforcing sex-specific adaptations.

## In a lone context, mice exhibit two foraging strategies

We first assessed mice foraging behavior and underlying neural mechanisms in a lone context. Female or male mice (N=62) were placed alone in a 50 × 50 cm multi-compartment environment and tracked continuously for 5 days and 4 nights using the Live Mouse Tracker (LMT^25^) system (**Figure 1a**). This setup includes a lever on one side and a food dispenser with a beam on the opposite side, enabling mice to obtain pellets by pressing the lever and consume them at any time (**Figure 1a**). Mice displayed nocturnal foraging activity, with lever presses (LP) and nose-pokes in the food dispenser occurring primarily during the dark cycle, as expected (**Figure 1b, left**). Over time, animals increased LPs and reduced nose-pokes, indicating learned association between LP and food retrieval (**Figure 1b, right and Figure S1a-c**). Accordingly, trajectory durations between LP and food dispenser showed a skewed distribution, with 54% below 6 seconds, which we defined as a complete sequence (**Figure 1c, left**). The percentage of complete sequences (number of complete sequences / numbers of LPs) increased over sessions, stabilizing at 35%, suggesting instrumental use of LPs (**Figure 1c, right**).

**Figure 1:**
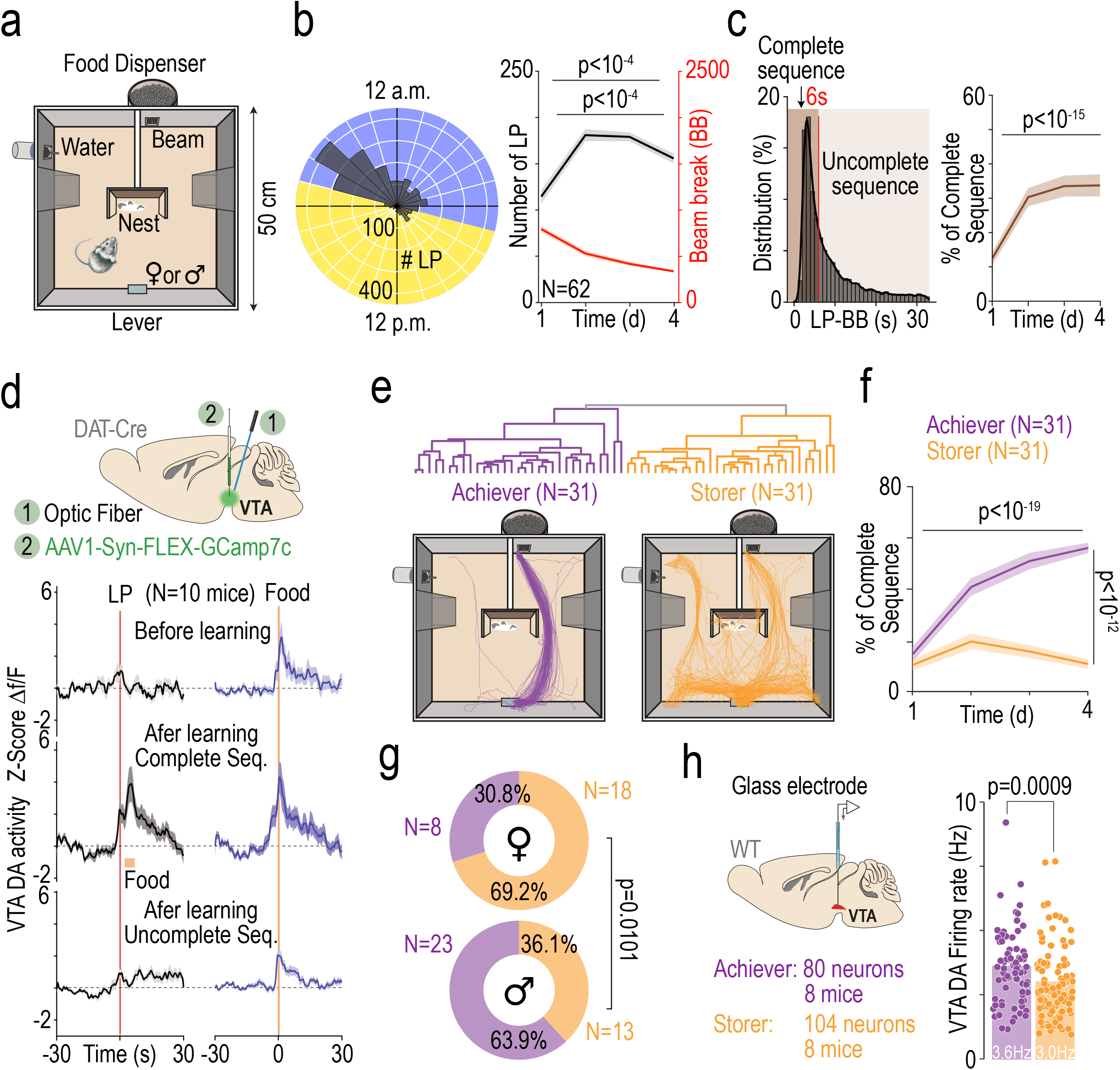
Different reward-seeking strategies emerged in a non-social semi-natural environment. **(a)** Mice were housed individually for 5 days in habitats where food was delivered via a lever-press (LP), positioned opposite to the food dispenser. **(b)** (Left) Cumulative polar LP histogram over 24 hours, showing peak activity during the dark cycle. (Right) Time course of LP and beam breaks over 4 nights. **(c)** (Left) Distribution of trajectory durations between LP and dispenser, with a 6 sec threshold distinguishing complete and incomplete (over 6 sec) sequences. (Right) % of complete sequences over time. **(d)** (Top) Schema of injections and implantations. (Bottom) VTA DA activity (fiber photometry) differs before and after learning (after day 3) and based on sequence completion, with activation at food delivery. **(e)** (Top) Clustering distinguished two food seeking strategies: “*Achievers*” and “*Storers*”. (Bottom) Examples of 6 sec trajectory after a LP, from two different mice. **(f)** Percentage of complete sequences over time by strategy. **(g)** Strategy distribution by sex. **(h)** (Left) In-vivo electrophysiology setup. (Right) VTA DA activity in anesthetized males at experiment end, depending on strategy.

To explore the neural bases of this association, we used *in vivo* fiber photometry in DAT-Cre mice expressing GCaMP7c in ventral tegmental area (VTA) dopaminergic (DA) neurons (**Figure 1d and Figure S1d**). A phasic increase in VTA DA neuron activity was observed at the time of food delivery, regardless of the completeness of sequence types or of learning progression, consistent with reward signaling (**Figure 1d**). In contrast, we found a specific phasic increase in VTA DA neuron activity at LP time, but only after the learning phase and for complete sequences, consistent with prediction of food reward upon LP^31^. These findings further suggests that LPs associated with complete sequences acquired an instrumental status during learning, reflected in reward prediction-like VTA DA activity.

We next analyzed individual decision-making strategies. Clustering analysis, based on the number of LPs and the percentage of complete sequences, identified two distinct groups of mice: *Achievers* (N=31) and *Storers* (N=31) (**Figure 1e**). While these groups displayed a similar number of LPs, their percentages of complete sequences were different (**Figure 1f, Figure Supp 1d**). Specifically, *Achievers* made a high proportion of complete sequences, whereas *Storers* often made several LPs before retrieving the food, indicating different foraging strategies. Moreover, their distribution differed between sexes, with a higher proportion of *Storers* in female mice than in males (**Figure 1g**). Spontaneous VTA DA neuron activity – recorded post-experiment in anesthetized males– showed higher firing frequency in *Achievers* than in *Storers* (**Figure 1h, Figure Supp 1e**), suggesting a link between VTA DA activity and foraging strategies^26,32,33^.

## Distinct sex-specific strategies emerge within groups

We next explored the foraging strategies of WT mice (N=84) in a social context, by housing triads of either all-male or all-female task-naive mice for 8 days and 7 nights (**Figure 2a, top**). Overall, LP activity mirrored that in the lone context, with higher nocturnal activity (**Figure 2a, bottom**). Clustering, based on the percentage of LPs and complete sequences, identified four distinct foraging strategies (**Figure 2b-c**): i) *Independents*, who pressed the lever sufficiently for self-feeding (i.e., 150 pellets per day) and thus showed minimal social impact, ii) *Workers*, who pressed the lever extensively, producing more food than they consumed, effectively subsidizing other group members, iii) *Scroungers*, who rarely pressed the lever and mainly relied on food produced by others, and (iv) *Storers*, who, as in the lone context, frequently pressed the lever but, contrarily to W*orkers, Independents, and Scroungers*, performed few complete sequences. These roles or profiles rapidly emerged, with distinct behaviors observable from day 1 (**Figure Supp 2a**) and maintained over time (**Figure Supp 2b)**. Indeed, within hours on day 1, *Workers* exhibited frequent LPs and complete sequences, while *Scroungers* displayed few LPs and complete sequences. Similarly, *Storers* began pressing early (within the first five hours) and frequently, while performing few complete sequences. On the first day, *Storers* experienced frequent food “theft” (i.e., the food they produced was consumed by others), but also engaged in theft themselves (**Figure Supp 2c-d**). These results indicate that role specialization and consistent strategies develop rapidly and are stable.

**Figure 2:**
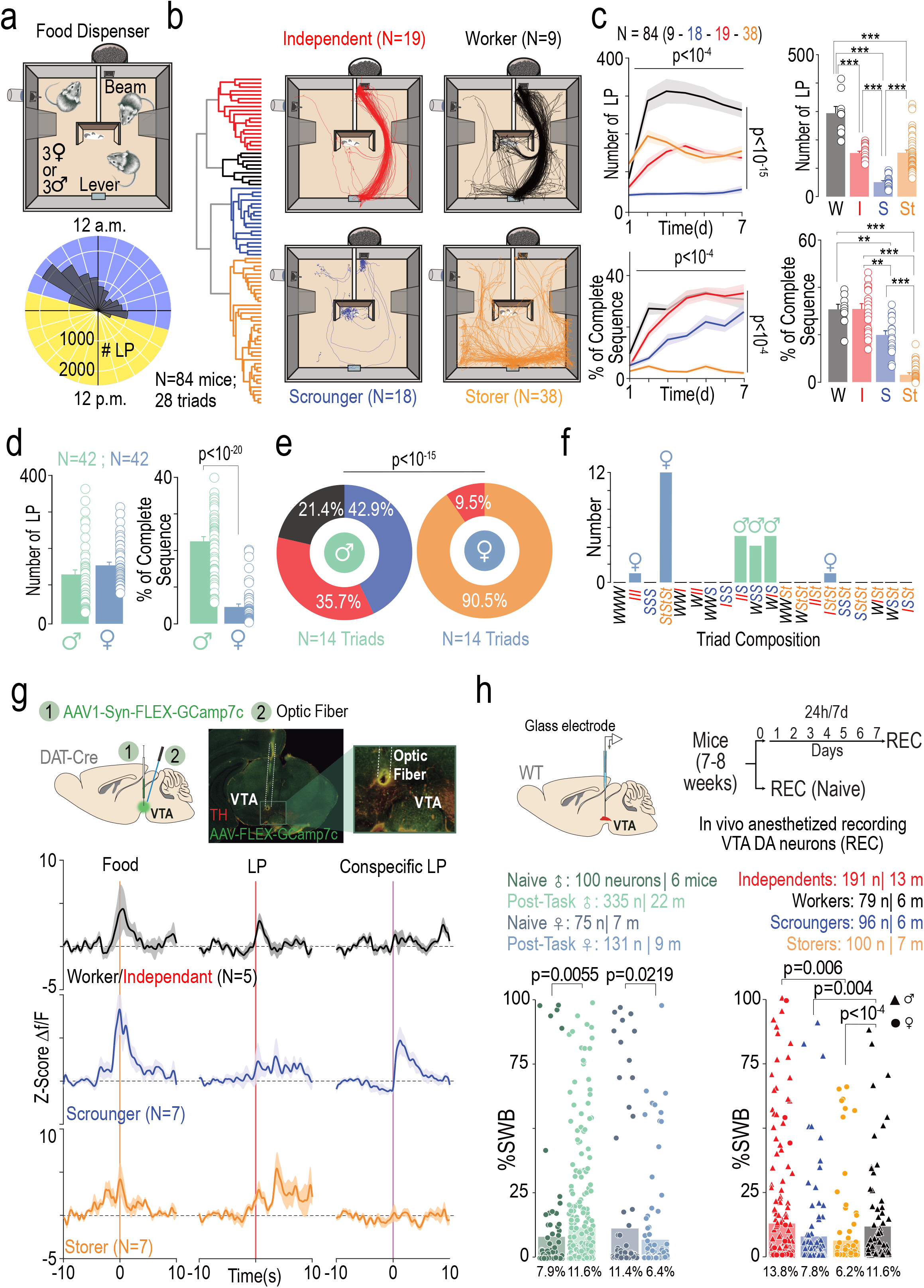
Distinct sex-specific strategies emerge within groups. **(a)** (Top) Triads of mice in the same environment as figure 1. (Bottom) Polar histogram of cumulative LP over 24 hours. **(b)** (Left) Clustering analysis identified four strategies *Independents, Workers, Scroungers*, and *Storers*. (Right) Example of a 6 sec trajectory after LP for each strategy, except for *Scroungers*, for which trajectory after Worker LPs is shown. **(c)** Time course of LPs (top left) and % of complete sequences (bottom left), with strategy-averaged values for the last two days (top right, bottom right). **(d)** Mean number of LPs and % of complete sequences by sex. **(e)** Strategy distribution by sex. **(f)** Triad composition **(g)** (Top) Schema of injection and implantation for fiber-photometry recording and example image. (Bottom) VTA DA activity varies at Food, LP, and conspecific LP depending on strategy. **(h)** (Top) In vivo electrophysiology setup and protocol. (Bottom) Firing patterns of VTA DA cells (percentage of spikes within burst - %SWB) in anaesthetized mice: comparison between naïve and experienced mice (left) and across strategies (right).

Sex-based differences were evident in both role types and their distribution within each cage. While males pressed the lever as often as females on average, they showed greater variability and more extreme behaviors **(Figure 2d, Supp 2e)**. Females also performed significantly fewer complete sequences **(Figure 2d, Supp 2e)**. Male triads (*n* = 14) typically comprised one or two *Scroungers* coexisting with *Workers* and *Independents*, but no *Storer* (**Figure 2e-f**), reflecting pronounced specialization. In contrast, female triads displayed remarkable uniformity in foraging strategies, with approximately 90% *Storers* (**Figure 2e-f**). This pronounced divergence between male and female triads suggests potential sex-based differences in neural, cognitive, and behavioral properties and/or social dynamics. Furthermore, the non-uniform distribution of profiles within triads **(Figure 2f)**—with certain triad compositions being absent, rare, or frequent—indicates that individual profiles are stabilized through dynamic social interactions. For instance, the emergence of three *Scroungers* appears unsustainable, suggesting that intrinsic social constraints govern the feasibility of specific role distributions.

To investigate the neural underpinnings of these behavioral strategies, we recorded VTA DA activity during the task, using fiber photometry in DAT-Cre mice (N=19) (**Figure 2g**). All groups showed increased VTA DA activity at food delivery, but *Workers* and *Independents* displayed a sharp rise during LP for complete sequences (**Figure 2g**), consistent with the use of LP as an instrumental action predicting reward. In contrast, *Scroungers and Storers* lacked this response, with *Scroungers* instead displaying heightened DA activity when conspecifics pressed the lever (**Figure 2g**), indicating that for scroungers, LPs of their peers, rather than their own, acquired the status of a reward predictor. This suggests a social alertness mechanism in *Scroungers* for tracking food produced by others. These distinct dopaminergic signatures cross-validates the behavioral classification (**Figure 2b)** and strengthen the link between observable strategies and their neural substrates. Finally, post-experiment recordings of VTA DA neurons revealed sex- and strategy-dependent differences (**Figure 2h, Figure Supp 3**): males in social contexts exhibited increased VTA DA bursting activity compared to unexposed males, whereas females showed the opposite. *Workers* and *Independents* had higher DA bursting activity than *Scroungers* and *Storers*. These results underscore the dynamic and sex-dependent interplay between DA signaling and behavioral roles, suggesting plastic neural adaptations to social contexts.

## Sex-specific decision making in social context accounts for behavioral specialization

To assess the mechanisms of behavioral specialization, we developed a reinforcement Q-learning model (*Full model*, see methods). In this model, one or three e-mice learned to find food in an environment **(Figure 3a)** with four spatial states (1-4) and two action states — lever-press (L) and food dispenser (D)— where lever presses increase (and consumption decreases) reward availability by one pellet. Over time, state transition values formed a spatial gradient leading to the rewarded location (D; **Figure 3a**). Specialization could arise from intra- or inter-sex individual parameter differences, social constraints on behavior (i.e., lever press, food availability), or both. We assessed how model parameter variations could account for behavior specialization in lone and social conditions. Learning rate (*α*) and temporal discount factor (*γ*) parameters had moderate effects, whereas inverse temperature (*β*), which controls the exploitation-exploration tradeoff (high-valued / random decisions), was critical (**Figure Supp 4a-c, e-g**). In the lone context, varying *β* (**Figure 3a**) recreated the overall behavioral pattern observed in lone mice (**Figure 3b**). Lower *β* favored exploration, leading to shallower value gradients (**Figure Supp 4c**), and less constrained trajectories, resulting in lower percentage of complete sequences (**Figure Supp 4b**), resembling *Storers* (**Figure 1f**). By contrast, higher *β* promoted steeper value gradients (**Figure 3a**) and higher percentage of complete sequences (**Figure Supp 4b**), akin to *Achiever*s (**Figure 1f**). Crucially, introducing intra- and inter-sex–specific *β* variability — where males exhibited higher *β* values than females (**Figure 3c)** — allowed the model to replicate the observed sex differences **(Figure 3b)**.

**Figure 3:**
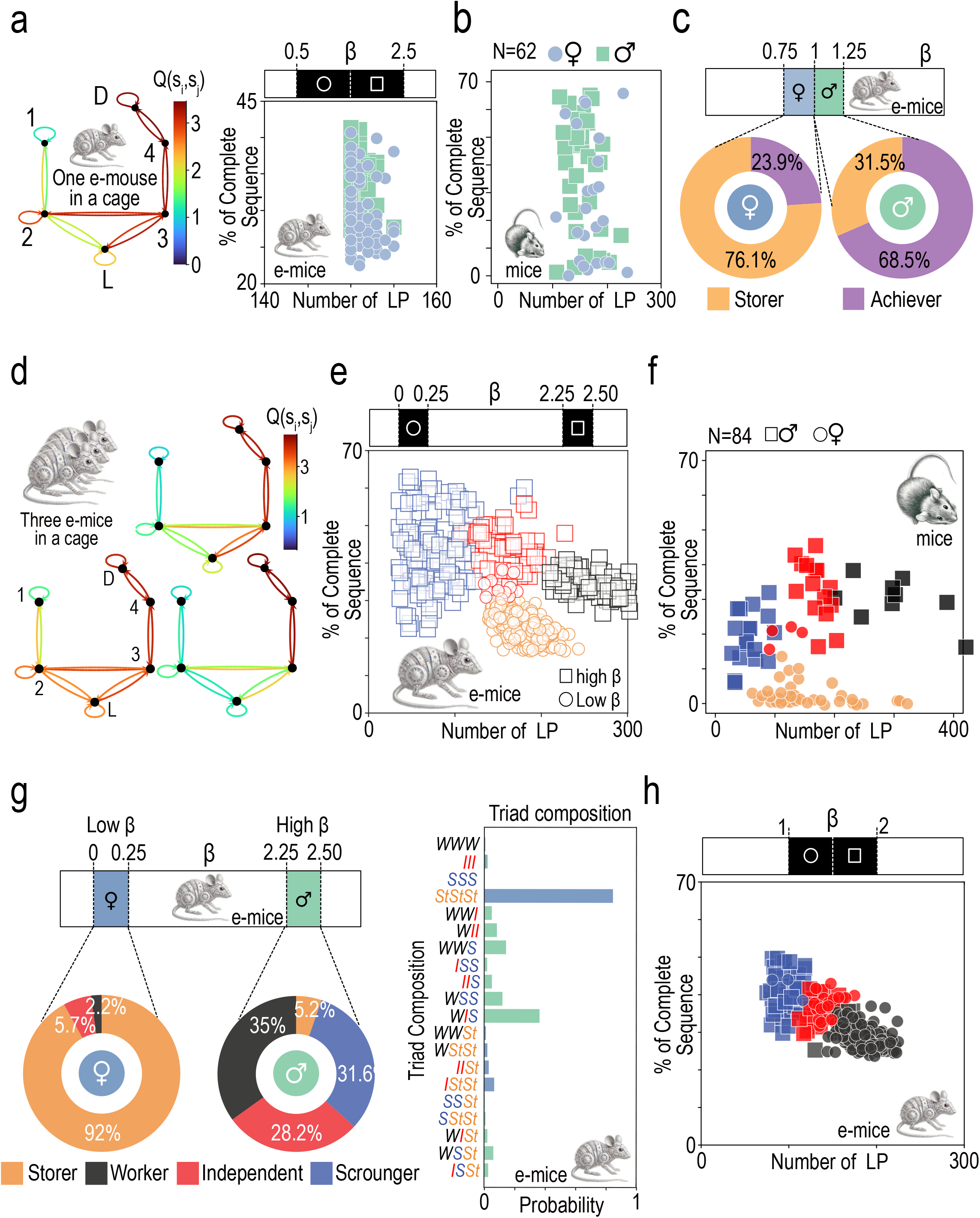
Reinforcement learning model and emergent strategies. (a) (Left) RL model (full model, methods) with one e-mouse, four states and two actions: lever press (L) and eat at dispenser (D). Q-values represent agent learning (e-mice). *β*=2. (Right) LP count vs. % of complete sequences for 200 e-mice with β uniformly distributed in [0.5, 2.5]. **(b)** Experimental LP vs. % of complete sequences for 62 mice, with males (empty squares) and females (filled circles). **(c)** Mean proportions of *Achievers* and *Storers* over repeated clustering (see methods) **(d)** Simulation as in (a), with three e-mice. **(e)** LP vs. % of complete sequences for 200 e-triads, clustered into *Independents* (red), *Workers* (black), *Scroungers* (blue), and *Storers* (yellow). **(f)** Experimental data for 84 mice, with color code for profile as in (e). **(g)** Strategy distribution and triad composition for 200 single-sex e-triads. **(h)** LP vs. % of complete sequences for 200 mixed e-triads (two females, one male) clustered into *Independents* (red), *Workers* (black), and *Scroungers* (blue).

Strikingly, even with identical *β*, e-triads (e-mice triads) with high *β*s (favoring exploitation; *β* ⪆ 1), displayed distinct gradients and behaviors (**Figure 3d**), whereas e-triads with low βs (favoring exploration; *β* ⪅ 1) displayed no specialization (**Figure Supp 4d**). A narrow high-*β* distribution produced large variability in percentage of complete sequences, with clustering unravelling *Worker, Independent* and *Scrounger* types, whereas (**Figure 3e**) a narrow low *β* distribution produced S*torer* e-triads. Thus, sex-specific *β* distributions produced patterns that closely matched those observed in male and female triads (**Figure 3f**). Moreover, these *β* distributions accounted for the distribution of behavioral types and triads (**Figure 3g**) found in mice.

In contrast, contiguous low and high *β* distributions with a larger variability led to specialized profiles without Storers (**Figure 3h**), as well as specific types and e-triads distributions (**Figure Supp 4h, i**), with high *β* individuals as *Scroungers* or *Independents* and low *β* as *Independents* or *Workers* (**Figure 3h**).

## Contingence and competition constrained behavioral specialization

We refined the causal mechanisms of social behavioral differentiation in a mathematically tractable reduced model (see methods) with two e-mice and two locations (L and D). Social differentiation arose through symmetry breaking of individual behaviors at a supercritical pitchfork bifurcation at *β*=*β*_Bifurc_ value (**Figure 4a**). Below *β*_Bifurc_ (e-females), the single fixed-point for Q-values toward D (*q*_*D*_) and residence probability at D (*pD*), lever press (*pLP*) and complete sequence (*p*_*PS*_) probabilities corresponded to *Storer* (**Figure 4a** and **Figure Supp 5a-c**). Above *β*_Bifurc_ (e-males), the two stable fixed points branches, correspond to *Workers* and *Scroungers* (bistability; **Figure 4a** and **Figure Supp 5a-c**).

**Figure 4:**
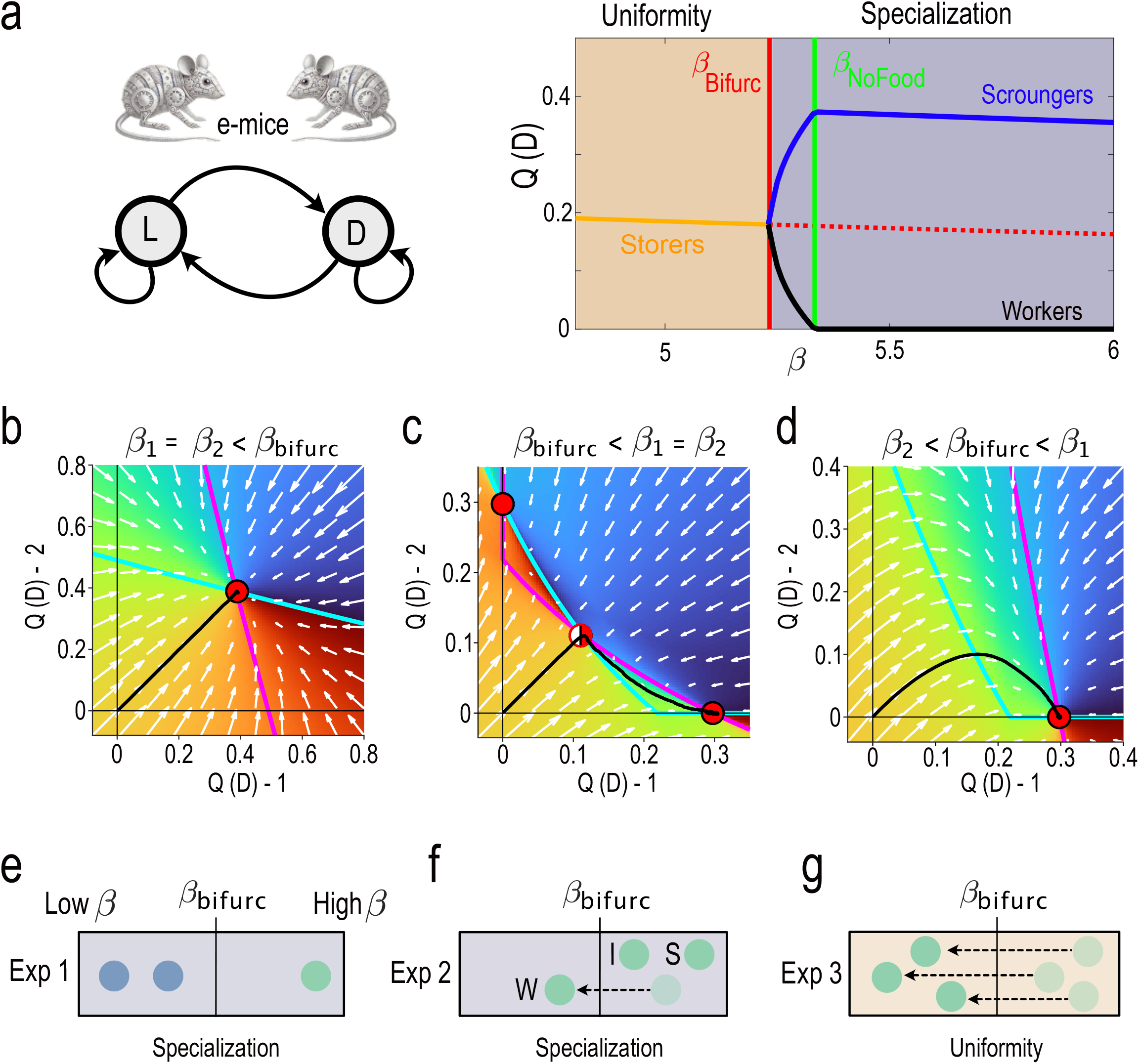
Behavioral specialization from competition-induced bifurcation. **(a)** Reduced model with two states (L: lever, D: dispenser) and two mice (1 and 2). (Right) Bifurcation diagram of the Q-value at D as a function of *β*, with two characteristic values, *β*_bifurc_ a bifurcation point and *β*_NoFood_ (see methods). (b) Similar low *β*s e-mice results in a single stable fixed point (uniform profiles) in the velocity landscape Black curve: simulated dynamics example. **(c)** Similar high *β* values yields two stable fixed points corresponding to a *Worker (here e-mouse #1)* and a *Scrounger*. **(d)** One high-*β* e-male and one low-*β* e-female yields a stable fixed point, with a male *Scrounger*. **(e-g)** Experimental design to test model predictions. **(e)** A triad with a high-*β* male and a low-*β* female pair to assess role specialization. **(f)** Lowering β in a male to test the likelihood of adopting *Worker* roles. **(g)** Reducing *β* in a male triad to evaluate the emergence of *Storer*-like dynamics analogous to female triads.

Qualitative analysis^34^ also revealed the causal origin underlying symmetry breaking and behavioral bistability for e-male (**Figure 4b-c** and **Figure Supp 5d-e**). Initially, contingency (i.e. individual random choices) determined which e-mouse first accessed the dispenser (D), increased its Q-Value (*q*_*D*_) and selection probability (*p*_*D*_), which both were then amplified through the positive feedback loop between Q-learning and softmax decision processes. This feedback led the e-mouse to predominantly occupy the dispenser area and become a *Scrounger*. The reduced food access for the other e-mouse of the dyad implied low reward and learning at D, and consequently, a higher presence probability at L (i.e., more lever presses). The other e-mouse thus became a *Worker*. Thereby, competition for shared resources is the causal factor driving behavioral specialization in male e-triads. Higher *β* intensified specialization, indicating that increased competition due to greater resource exploitation promoted differentiation in males. In contrast, e-female maintained uniform behaviors, attributed to lower levels of exploitation and resource competition.

Behavioral specialization thus arises from several intricate mechanisms. Sex-specific difference in *β* determines whether e-mice become *Storers* (females) or specialize into distinct behavioral profiles (males). In males, specialization arises from both random contingency and higher *β* values, with larger *β* e-mice being more likely to become *Scroungers*. Therefore, the model highlights *β* as a key determinant of behavioral specialization and social organization, and differences in exploitation between males and females as the critical factor underlying sex-specific social structures. Additionally, the model predicts that pairing high-*β* with low-*β* individuals should lead to a *Scrounger-Worker* dynamics, where the more exploitative individual becomes *Scrounger*, while the less exploitative one becomes the Worker **(Figure 4d and 3h)**. Three experiments can empirically test these predictions. First, introducing a single high-*β* male with a low-*β* female pair should induce specialization in females, shifting them toward *Workers* or *Independents* (**Figure 4e**), and damper the competition between males, resulting in fewer *Scrounger*s. Second, lowering *β* in one male within a triad should increase its likelihood of becoming a *Worker* **(Figure 4f)**. Finally, reducing *β* in all three males should produce a *Storer*-like dynamics, as seen in female triads **(Figure 4g)**. We next experimentally tested these hypotheses to validate our model’s predictions.

## Testing model predictions with mixed-sex triads: The influence of low-*β* females on specialization

We experimentally tested the first of these predictions **(Figure 4e)** by forming mixed-sex triads composed of one male and two females (**Figure 5a**, Methods). Males performed fewer LPs than females, but unlike in sex-matched triads, there was no difference in the percentage of complete sequences between sexes (**Figure 2d**). Clustering analysis identified three profiles — *Scroungers, Workers* and *Independents* (**Figure 5b**) — mirroring those seen in male-only triads. Females neither adopted the *Storer* profile (due to increased competition) nor the *Scrounger* profile (due to their lower *β*s). Furthermore, in nearly half of the triads (5/12 triads), no *Scrounger* mouse emerged, validating our first prediction: incorporating low-β individuals (females) reduced the competition driving specialization, instead fostering more cooperative dynamics. The emergence of an individual’s role is thus not solely driven by its own traits, but by the dynamics induced by trait distribution within the group.

**Figure 5:**
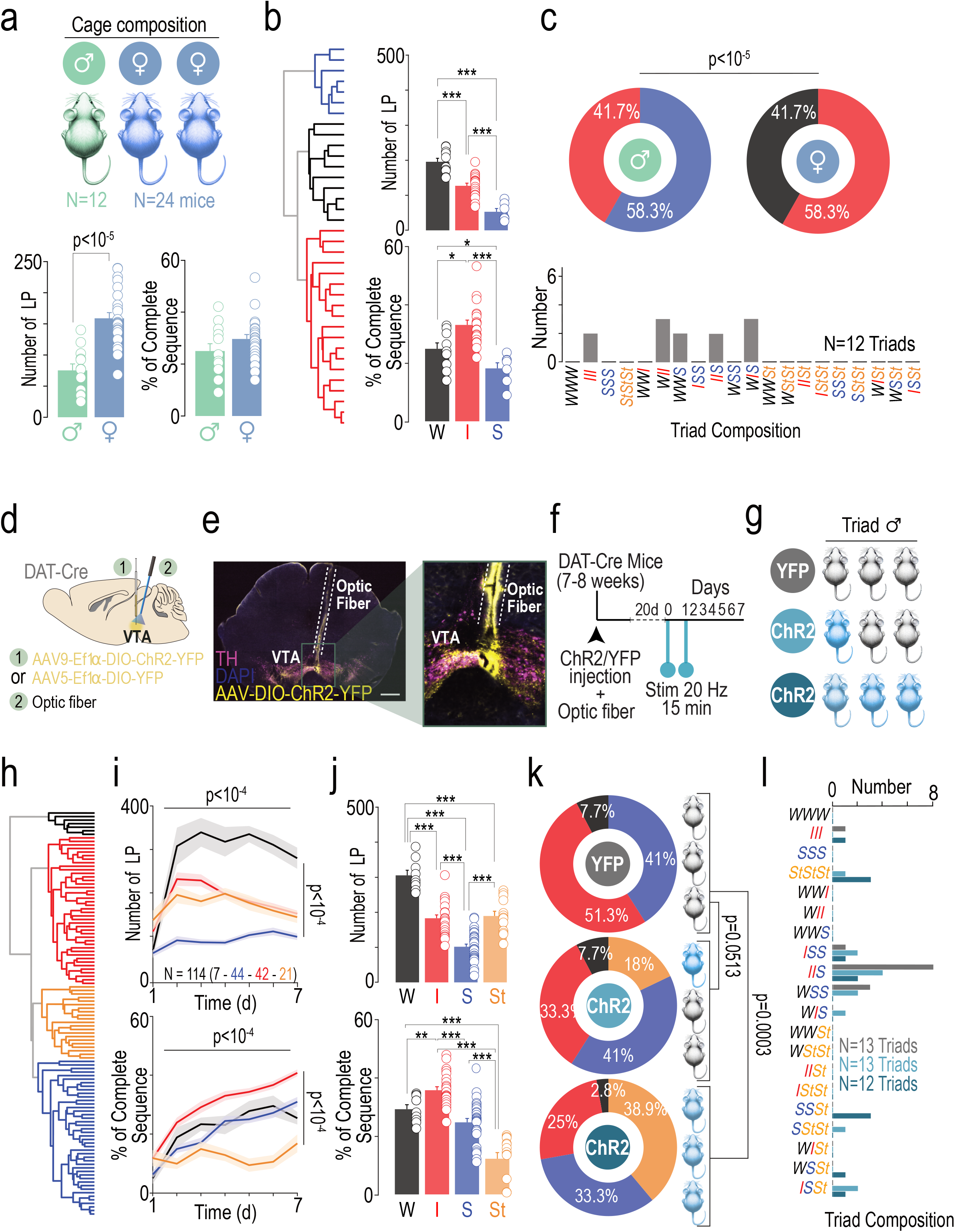
Testing model predictions and exploring the role of VTA dopaminergic activity in behavioral specialization. **(a)** (Top) Mixed-sex triads (1 male and 2 females). (Down) Mean LP and % of complete sequences for males (green N=12) and females (blue N=24). **(b)** (Left) Clustering identified three strategies: *Scroungers, Workers*, and *Independents*. (Right) Number of LPs and % of complete sequences for the last two days per strategy. **(c)** (Top) Strategy distribution by sex. (Bottom) Triads composition. **(d)** Experimental design for VTA DA manipulation using ChR2 in male DAT-Cre mice. **(e)** Example image of viral infection and fiber implantation. **(f)** Experimental setup: VTA DA stimulation applied 24 hours and 30 minutes before microsociety experiment. **(g)** Triad compositions: (i) all YFP controls, (ii) two YFP with one ChR2-stimulated mouse, (iii) all ChR2-stimulated. **(h)** Clustering in optogenetically manipulated male triads identified four strategies: *Workers, Independents, Scroungers*, and *Storers*. **(i)** Number of LPs (top) and % of complete sequences (bottom) over time by strategy. **(j)** Mean LP (top) and % of complete sequences (bottom) for the last 2 days. **(k)** Strategy distributions across triad types: (i) all YFP control mice, (ii) one ChR2-stimulated mouse with two YFP controls, and (iii) all ChR2-stimulated mice. **(l)** Triad composition.

## Artificially increasing VTA DA neuron activity influences role specialization in male microsocieties

VTA DA activity has been linked to exploration-exploitation parameters, further supporting its involvement in shaping role-specific behaviors ^24,33,35,36^. Accordingly, we observed variations in tonic VTA DA neuron activity at the end of the experiment across behavioral roles and sexes. While our model does not explicitly predict a direct relationship between *β* and VTA DA firing rate, we tested whether modulating DA activity could impact specialization dynamics **(Figure 4f, g)**. We selectively increased basal VTA DA neuron activity in male DAT-Cre mice^37^ using excitatory channelrhodopsin (ChR2) (**Figure 5d, e, and Figure Supp 6a-d**). Mice received 15 minutes of photostimulation 24 hours before, and again 30 minutes before the start of the microsociety experiment (**Figure 5f**). Electrophysiological recordings confirmed that this stimulation reliably increased VTA DA neuron activity for at least 6 hours post-stimulation (**Figure Supp 6a-d**). This allowed us to assess whether altering one (**Figure 4f**) or all (**Figure 4g**) individual’s DA activity affected role distribution.

We thus compared three types of male triads: (i) three YFP control mice (n=13 triads), (ii) one ChR2-stimulated mouse and two YFP controls (n=13 triads), and (iii) three ChR2-stimulated mice (n=12 triads). Behavioral clustering identified four strategies – *Workers, Independents, Scroungers*, and *Storers* (**Figure 5h**) – whose behavior was consistent with our previous observations (**Figure 5i-j ; Figure 2b,c**). Surprisingly, *Storer* strategy emerged in photostimulated male triads, shifting the overall role distribution compared to YFP controls (**Figure 5k**; chi-squared test, p = 0.051). Importantly, the *Storer* was not necessarily the stimulated individual, emphasizing the role of group dynamics in shaping behavioral outcomes **(Figure Supp 6e)**. This suggests that enhancing VTA DA activity prior to the social experiment leads to a lower β, facilitating the emergence of Storers by diminishing exploitative strategies. This effect was further confirmed in triads where all three mice were ChR2-stimulated, in which *Storer* frequency was increased (**Figure 5k**; chi-squared test, p = 0.0003). These findings demonstrate that while increasing VTA DA activity did not disrupt existing roles, it facilitated a female-like *Storer* profile in male triads. This highlights the complex interplay between VTA DA neurophysiology and the social environment in shaping individual strategies, providing a neural basis for the adaptive specialization observed within these microsocieties.

## Discussion

In this study, we highlight the importance of exploring social organization through the lens of division of labor to gain deeper insights into behavioral specialization in animal microsocieties. Traditional accounts of behavioral specialization depict fixed social roles as evolutionarily stable strategies^20,38^, but our findings challenge this static view by instead revealing a dynamic, learning-driven processes ^21,23^. Our results demonstrate that social norms, conceptualized as implicit rules governing group behavior, emerge rapidly from animals’ coexistence and dynamical structure, guiding both individual roles and collective dynamics.

A key contribution here is the demonstration that an individual cognitive property, i.e. the exploration-exploitation tradeoff, constraints the emergent social structure. Using reinforcement-learning models and experimental data, we show that variability in neural mechanisms, particularly dopaminergic signaling in the VTA, shapes how behavioral roles evolve. This neurophysiological foundation emphasizes that behavioral specialization is not merely an adaptive response to external social demands, but arises from a permanent causal interaction between intrinsic neural properties and social interactions, through a feedback loop: social contexts constraints individual’s neural dopaminergic levels, which in turn reinforce specific behavioral strategies, eventually stabilizing social structures over time.

A striking aspect of our findings is the pronounced sex-specific divergence in social organization. Male groups exhibited distinct and stable divisions of labor, including roles such as *Workers, Scroungers* and *Independents*, whereas female groups displayed remarkable behavioral uniformity, predominantly behaving as *Storers*. These differences highlight how sex-based cognitive and neurophysiological differences interact with social environments to shape collective dynamics. For example, male-specific dopaminergic activity patterns, coupled with higher exploitation behavior (high *β*), promote competition and role specialization. Conversely, females’ explorations tendency (low *β*) appears minimize competition and favor behavioral uniformity. These findings resonate with observations that males and females exhibit distinct responses to social competition and resource access ^39^. Males are more prone to develop divergent roles under competitive pressures, while females tend to maintain consistent strategies, highlighting sex-specific adaptations in response to social dynamics.

The role of early-life contingency in shaping behavioral and social outcomes warrants discussion. It has been proposed that early-life experiences and chance events can magnify small initial differences between individuals, leading to diverging trajectories^24,29,39–41^. In competitive social environments, such as those often encountered by male mice, these feedback loops amplify disparities and drive the emergence of distinct behavioral roles. By contrast, female mice may experience reduced competitive pressures, allowing for more equitable outcomes within their groups. In the model, *β* distributions required to reproduce experimental behaviors were compatible with the hypothesis that the presence of other males or other females in the unisex triads increases or decreases exploitation, respectively, possibly due to social effects on the internal state of the individual ^3,42,43^. This contingency-based framework emphasizes the importance of considering both stochastic aspects (e.g., unpredictable group composition or initial random reward discovery) and deterministic factors (e.g., one animal becoming a *Scrounger* favors others to become *Workers*) in shaping sex-specific behaviors and social structures.

Our findings demonstrate that dopaminergic signaling plays a central role in behavioral specialization, with *Workers* and *Independents* showing VTA activity linked to reward prediction, while *Scroungers* respond more to conspecific lever presses, suggesting social alertness. These distinct dopaminergic profiles reinforce the idea that neural plasticity underlies the emergence of specialized roles within social groups. Furthermore, variations in tonic VTA DA activity, shaped by both intrinsic predispositions and social experiences, likely contribute to the stabilization of individual strategies over time. Future research should determine whether these neural adaptations persist across successive different social settings or revert when individuals are removed from their original groups.

In conclusion, this work provides a comprehensive analysis of how individual and social factors interact to shape behavioral specialization in animal microsocieties. By bridging neurophysiology, behavioral science, and computational modeling, our findings offer a nuanced understanding of the dynamical interplay between individual cognition and collective behavior, with broad implications for the study of social systems across taxa.

## Methods

### Animals

Males and females wild-type (WT) C57BL/6J mice (8–12 weeks old, Janvier Labs, France) or DAT-iCre mice (C57BL/6J background, BAC-DAT-iCre, from François Tronche^44^) were housed in groups in an animal facility and maintained on a 12-hour light–dark cycle (7:00 a.m.–7:00 p.m.), at an average temperature of 22°C (22 ± 2°C) and ∼50% humidity. All the animals lived in triads in the animal facility before entering in the task for 8 days with the same triad for the same-sex experiments. In lone condition, the mice were housed individually in the cage for 5 days only, to prevent side effect of long-term social isolation.

All experiments and procedures were performed in accordance with European Commission directives 219/1990, 220/1990 and 2010/63, and approved by the ESPCI and the ethical committee #059 under APAFIS #34335-2021121318085835.

### Behavior

Mice were placed either individually or in groups of three (triads) in a 50 × 50 cm large environment and continuously tracked using the LMT system^25^, allowing to keep the identity of each animal over extended periods. The mice in groups (3 males, 3 females or 2 females / 1 male) stayed for 8 days and 7 nights in the environment, while mice placed alone in the cage stayed only 5 days and 4 nights.

The cage was composed of different zones freely accessible by all the mice at any moment. A lever was placed on one side of the environment, while a food dispenser with a magazine were located at the opposite side. Every time the lever is pressed, a small food pellet of 20 mg (TestDiet 5TUL/1811142 purified, Bio-Concept) is delivered at the magazine and the lever becomes inactive for 5 seconds. A nose-poke in the magazine leads to a beam break that is a proxy of the animal’s consumption of the pellet. Outside the 5 seconds delay between each lever-press, all the mice can press at any moment and consume the pellet at any time following its release. The mice were checked and weighted every day to keep track of the good health of each individual. No mouse was removed from the task either for weight loss or for aggression.

### Live Mouse-Tracker system

This setup, in our paradigm, provided a controlled context for examining how individual mice learn and adapt to the task, either in isolation or within a standard social setting, allowing comparisons between lone and group behaviors. We employ comprehensive 24/7 data collection within the experimental environment, a method that provides rich longitudinal datasets. The data collected fall into two main categories: discrete events and continuous video streams. Discrete events are automatically captured by sensors integrated into the environment, including infrared beams at the lever and nose poke, and food dispenser activations (MedAssociates). Each event is tagged with the corresponding sensor number, timestamp, and the identity of the animal involved. These events are logged into a structured database, enabling detailed analysis of individual and group-level behavioral patterns over time. In addition to discrete events, we record continuous 24/7 video streams, which provide a complete visual record of animal’s behavior within the environment. To manage the substantial storage demands of such video data, we implement real-time processing techniques to extract essential features. This includes tracking each animal’s center of mass and body contours, allowing us to quantify locomotion, spatial organization, and interactions. These extracted metrics are stored in a compressed format for subsequent analysis, ensuring that critical information is retained without overwhelming storage capacity. This dual approach to data collection provides a nuanced understanding of behavior, capturing both fine-grained event data and broader spatial and social dynamics.

Every day the LMT and MedAssociates softwares were relaunched to prevent huge file weight. The number of lever-presses, the number of nosepokes and the number of complete sequences were extracted directly from the database using PyCharm (Python) and MySQL. We defined two specific zones around the lever and the magazine to be able to know which animal was located there during a lever press or a nose poke. We then determined which TTL was associated with which animal in the zone. For each day of analysis, it was not possible to associate a little percentage of trials (< 5%) with specific mice since it happened that several mice were present at the same time either at the lever or at the magazine. Once the association made between the position of the animal at a specific timestamp and the time of the TTL, we obtained the number of lever-presses and nose pokes for each animal. For complete sequences, we quantified the number of times that the mouse which had pressed the lever was the first arrived at the subsequent nose poke in less than 6 seconds, otherwise the sequence was not counted (typically if a mouse was already at the magazine waiting for the food for example).

### Behavioral analyses

After extraction of the data from the database using PyCharm and MySQL, MATLAB was used to perform more detailed analysis. The percentage (%) of complete sequences was calculated as the number of complete sequences divided by the number of lever presses.

The clustering analysis was made using “linkage” followed by “dendrogram” functions on MATLAB. We normalized all the data for all the mice as followed:

- For the lone context: we used % of complete sequences and the ratio lever-press on nose poke for each mouse. We then pooled all the mice of all the cages, males and females, and we performed the clustering on the last two days of the experiments.
- For the social context: we used % of complete sequences and % of lever presses in the triad (number of lever presses of one mouse divided by the total number of lever presses of the triad). We pooled all the mice together, males and females for figure 2, and all the virus conditions for figure 5, independently of the triad. We performed the clustering analysis on the last five days.

### In vivo electrophysiology

Mice were deeply anaesthetized with isoflurane delivered continuously (3% induction, 1-2% maintenance; TEMSega). The scalp was opened and a hole was drilled in the skull above the location of the VTA. Intravenous administration of saline or nicotine (30µg/kg) was carried out through a catheter (30G needle connected to polyethylene tubing PE10) connected to a Hamilton syringe, into the saphenous vein of the animal. Extracellular recording electrodes were constructed from 1.5 mm outer diameter / 1.17 mm inner diameter borosilicate glass tubing (Harvard Apparatus) using a vertical electrode puller (Narishige). The tip was broken straight and cleaned under microscopic control to obtain a diameter of about 1 µm. The electrodes were filled with a 0.5% NaCl solution containing 1.5% of neurobiotin^®^ tracer (VECTOR laboratories) yielding impedances of 6-9 MΩ. Electrical signals were amplified by a high-impedance amplifier (Axon Instruments) and monitored audibly through an audio monitor (A.M. Systems Inc.). The signal was digitized, sampled at 25 kHz, and recorded on a computer using Spike2 software (Cambridge Electronic Design) for later analysis. The electrophysiological activity was sampled in the central region of the VTA (coordinates: between 3.1 to 4 mm posterior to bregma, 0.3 to 0.7 mm lateral to midline, and 4 to 4.8 mm below brain surface). Individual electrode tracks were separated from one another by at least 0.1 mm in the horizontal plane. Spontaneously active DA neurons were identified based on previously established electrophysiological criteria^45^. After recording, a subset of nicotine-responsive cells were labelled by electroporation of their membrane: successive currents squares were applied until the membrane breakage, to fill the cell soma with neurobiotin contained into the glass pipet^46^. To be able to establish correspondence between neurons responses and their location in the VTA, we labeled one type of response per mouse: solely activated neurons or solely inhibited neurons, with a limited number of cells per brain (1 to 4 neurons maximum, 2 per hemisphere), always with the same concern of localization of neurons in the VTA.

### Stereotaxic surgeries

For fiber photometry experiments: injection of AAV1-Syn-FLEX-GCamp7c (Titer ≥ 9.4×10^12^ vg.mL^-1^, Addgene, Watertown, MA, USA) was performed in 6-8 week-old DAT-Cre mice. Mice were anesthetized with a mixture of oxygen (1L/min) and isoflurane 2% (TEMSega, Pessac, France). After shaving the skin, mice were placed in a stereotaxic frame (David Kopf, Tujunga, USA), the skin was disinfected and locally anesthetized with 70 μL of lurocaine (0.5 %). Craniotomies were then performed, and unilateral injections were performed over the VTA at AP: -3.20 mm ML: ± 0.5 mm from the bregma and DV: −4.20 ± 0.05 mm from the brain surface. Injections were carried out with a glass micropipette (Drummond Scientific Company, Broomall, PA), at a 100 nL/min rate for a total volume of 300 nL in the VTA. At the end of the surgery, 100 μL of buprenorphine (0.1mg./kg) was injected subcutaneously. Unilateral optic fiber (200 µm core, NA= 0.39, ThorLabs) implantations were performed 2 – 3 weeks later above VTA with the same coordinates, and dental acrylic (SuperBond, Sun medical) was used for fixing the ferrule to the skull.

For optogenetic experiments: AAV5-Ef1α-DIO-ChR2(H134R)-EYFP (Titer ≥ 4.7×10^12^ vg.mL^-1^, VVF Zürich, Switzerland) or AAV5-Ef1α-DIO-EYFP (Titer ≥ 5.4×10^12^ vg.mL^-1^, VVF Zürich, Switzerland) were injected in 6-8 week-old DAT-Cre mice. Mice were anesthetized and disinfected as previously described. The animals were placed in a stereotactic frame and bilateral injections were performed in the VTA at the same coordinates than above, 300 nL per side, using a glass micropipette. After 2 – 3 weeks, one optic fiber (200 µm core, NA= 0.39, ThorLabs) was implanted unilaterally above the VTA with 10° angle (AP: - 3.20 mm ML: ±0.9 mm from the bregma and DV: −3.95 mm from the brain surface), and fixed to the skull using dental acrylic.

After all the surgeries, mice were placed alone in a heated cage for one hour recovery, and were subsequently checked every day. The behavioral experiments started at least 1 week after the surgery. Injections and implantations were all confirmed with post-hoc immunochemistry.

### Immunohistochemistry

Following euthanasia, brains were rapidly extracted and fixated in a solution of 4% paraformaldehyde for a minimum of three days at 4°C. Subsequently, serial sections of 60 micrometers in thickness were obtained from the midbrain using a vibratome.

For immunostaining experiments (TH-staining), free-floating brain sections were initially incubated for one hour at 4°C in a phosphate-buffered saline (PBS) blocking solution containing 3% bovine serum albumin (BSA) by volume/volume ratio (Sigma; A4503) and supplemented with 0.2% Triton X-100. This was followed by overnight incubation at 4°C with a primary antibody solution containing a mouse monoclonal anti-tyrosine hydroxylase antibody (anti-TH, Sigma, T1299) diluted 1:500 in PBS supplemented with 1.5% BSA and 0.2% Triton X-100.

The following day, sections were thoroughly rinsed with PBS and then incubated for three hours at room temperature (22-25°C) with the secondary antibody Cy3-conjugated goat anti-mouse (Jackson ImmunoResearch, 715-165-150) diluted at 1:500 in a solution of 1.5% BSA in PBS. After undergoing three additional rinses with PBS, the slices were mounted using Prolong Gold Antifade Reagent with DAPI (Invitrogen, P36930). Microscopic analysis was performed using a Zeiss epi-fluorescent microscope, and image capture was performed by a camera. To preserve the raw data, images were captured in grayscale format using the Zeiss software (ZEN microscopy software). Following acquisition, false coloring was applied using ImageJ to enhance visualization of the immunofluorescence signal.

### Fiber photometry

Mice injected with AAV1-Syn-FLEX-GCamp7c and implanted in the VTA underwent fiber photometry experiments in the lone or social context. A Doric Lenses fiber photometry system was employed to record fluorescence signals reflecting dopamine neuron activity in the VTA. The system utilized a fiber photometry console linked to an LED driver, controlling a connectorized blue light-emitting LED (465 nm) in Lock-in mode at 220.537 Hz. This signal was then transmitted through an optic patch cord to the Mini Cube (FMC4_AE(405)_E(460-490)_F(500-550)_S). An optical fiber, connected to the Mini Cube’s sample port and the animal’s implanted fiber via a zirconia sleeve, served as the conduit for both light stimulation and recorded fluorescence. The received light signal was converted to electrical signals by a photoreceiver using the AC Low setting, before being transmitted through another optic patch cord to the Mini Cube via a dedicated fiber optic adaptor. Finally, the Doric Neuroscience Studio software acquired the signal at a sampling rate of 12.0 kHz and applied a low-pass filter with a 12.0 Hz cutoff frequency. For lone condition, the animals were recorded between one and two hours at the beginning of the dark cycle, when the mice were active, during the first day and the last two last of the experiment.

For social condition, the animals were also recorded at the beginning of the dark-cycle, one after the other, between one and two hours each.

### Analysis of fiber photometry recordings

We first subtracted a biexponential detrend to our fluorescence signal in order to eliminate the slow decay of the signal due to sensor photobleaching, and then added on offset equal to the signal mean before detrending. The ΔF/F ratio was calculated as follow:

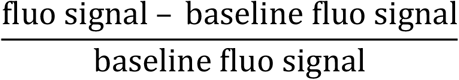

The analysis of VTA DA calcium variations and its correlation with behavior relied on the construction of peri-event time histograms, which were obtained by aligning and centering specific behavioral events. The identification of these events was done with TTLs received on the fiber photometry console at lever press or nose-poke events. The temporal resolution of the recordings was determined by the bin size chosen for analysis. We used a bin size of 100 ms to achieve a more precise representation of the calcium variations over time. Subsequent analyses were conducted on normalized activity data. To obtain normalized calcium fluorescence variations, a specific formula was applied: 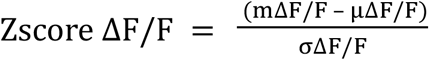, where Zscore ΔF/F represents the normalized calcium activity for a specific time bin, mΔF/F represents the average calcium signal variations within that specific time bin, µΔF/F represents the mean calcium signal variations across the baseline activity a few seconds before each events (5 seconds), and σΔF/F represents the standard deviation of the calcium signal variations across the baseline activity a few seconds before each events.

Following normalization, a smoothing technique was employed on the data. This involved convolution with a Gaussian kernel using a sliding window encompassing 100 bins. The “gausswin” function within the MATLAB software was utilized to achieve this smoothing process.

### Optogenetic experiments

In males DAT-Cre mice injected with AAV5-Ef1α-DIO-ChR2(H134R)-EYFP or AAV5-Ef1α-DIO-EYFP, optical stimulations were performed with an ultra-high-power LED (470 nm, Prizmatix) coupled to a patchcord (500 µm core, NA = 0.5, Prizmatix) with an output intensity of 5 – 10 mW. We used a 20 Hz stimulation protocol of 5 ms light pulse for 15 minutes 24 hours and 30 minutes prior to the social task or in vivo electrophysiological recording.

### Modeling

#### Building a behavioral model of e-mouse lone and social behaviors

The environment of experiments was modeled as 6 states (Figure 3a, d, rooms 1-4, and lever and dispenser positions). The number and sex of agents (“e-mice”) present in the environment was varied, with either 1) one (male or female) e-mouse in the lone experiment, 2) three male or female e-mice in social experiments, or 3) 1 male and 2 females in the mixed-box experiment. State transition occurred at each time step, with transitions probabilities determined by a softmax based on *Q* values of all accessible *k* states from the current state *s*_*i*_:

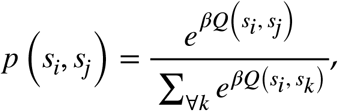

with *β*, the inverse temperature parameter, favoring either exploration (i.e., random choices; lower *β*s) or exploitation of previous learning (higher *β*s). *Q* learning occurred upon each transition from a departure state *s*_*i*_ to an accessible arriving state *s*_*j*_, as

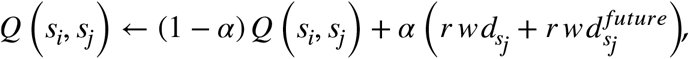

with α the learning rate, 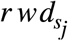 the reward obtained at the arriving state *s*_*j*_ and 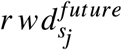 the maximal expected reward obtainable in states accessible from the arriving state *s*_*j*_:

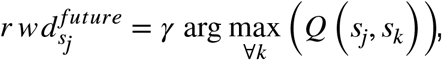

*γ* being a temporal discount factor. Furthermore, e-mice encountered satiety, which scaled action probabilities, the learning rate, and fatigue, which affected both pressing and eating. Satiety and fatigue were used to scale an action pace in the simulation that was consistent with the experimental measures. The lever could not be pressed for 3-time steps after each press (i.e., to mimic the 5 sec lever unavailability in experiments). The number of lever presses and the percentage of complete sequences depicted in Figure 3a, e and h were used to categorize e-mice and e-mice triads, as in experiments (Figure 3c, e, g, h, i). Complete modeling information regarding the full model is given in Supplementary Methods.

#### Reduced model of social interactions

We built a reduced theoretical model (hereafter termed the *reduced model*) to assess, within a mathematically tractable framework, the causal mechanisms whereby specialized behaviors emerge under social interactions. To do so, we derived the reduced model from the reinforcement-learning one, based on a continuous time version of *Q* dynamics (i.e., ordinary differential equations, ODEs). In these ODEs, learning and behavioral dynamics operated at a slower time scale – compared to that of individual choices in the full model – so that actions (i.e., state transitions, pressings, eatings) were described probabilistically. In this framework, we performed qualitative analysis of ODEs to determine the number and stability of fixed points of learning and behavioral (state) variables as a function of parameters (with as focus on *β*, which is essential in setting social interactions in the full model, see Results). Moreover, to reduce dimensionality for better tractability, we considered a simpler setup, where the environment contains only two positions for e-mice (the lever, *L*, and the food dispenser,*D*), only two e-mice (which allowed us to assess social interactions), and we did not consider fatigue or satiety. Reduced model ODEs could be expressed under a tractable form:

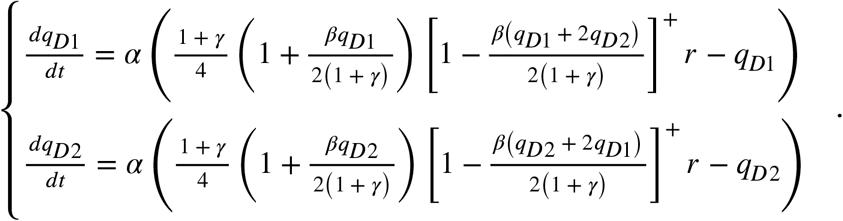

with *q*_*XYi*_ being the *q* value of *X* → *Y* transition and *r* the unit reward per eating. This system admitted one or several fixed points 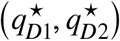, which number and nature depended on parameters (see the bifurcation schema and phase portraits, Figure 4a). We focused on the dependence of 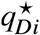 steady-states of individual e-mice on *β*, i.e., 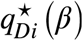, denoted *branches*. As in the full model, the value of *β* separated distinct regimes of social interaction between e-mice. We found that for *β* values below a critical value 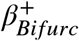, only one stable fixed point, 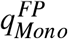, existed, with both e-mice admitting a similar *q* steady-state. Above 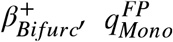 became a saddle node. Also, two other symmetric stable fixed points appeared,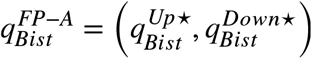 and 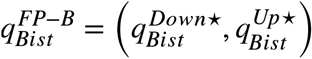,The fixed point 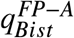 corresponded to the case in which e-mouse #1 displayed a higher *q* value at dispenser 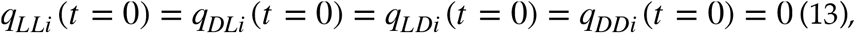, and, as a consequence, a higher residence probability 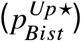 at dispenser, and lower probabilities of lever presses 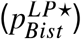 and direct sequences 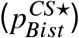, compared to e-mouse #2 (see below). In other words, 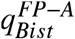 was a fixed point at which e-mouse #1 behaved as a scrounger and e-mouse #2 as a worker, while behaviors were inverted at 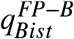. In this context, the 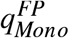 saddle node branch could be interpreted in this reduced model as corresponding to independent e-mice found in both social male boxes in experiments and in the full model (starting from (0,0) with small levels of noise yields animals to this saddle branche, see below). For *β* values above a second critical value 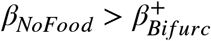, the probability of available food at the dispenser became null for the worker e-mouse (see detail of the analysis in Supplementary Methods), the system wrote differently and stable branches were continued by two branches 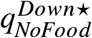 and 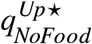. The main results of the qualitative analysis of the system are recapitulated in Table 1. Full information on the reduced model derivation and analysis in Supplementary Methods.

**Table 1:**
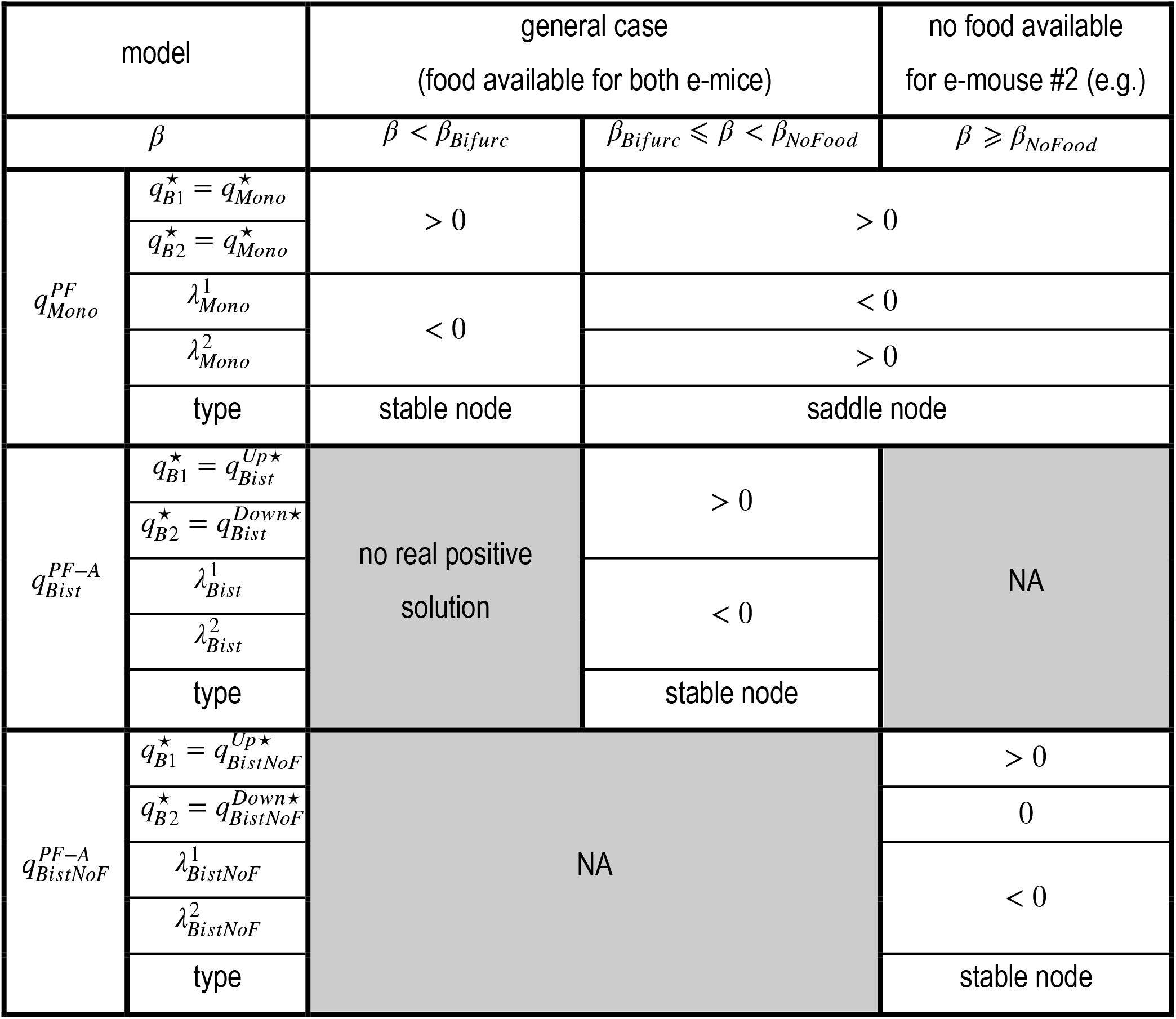
Model.

**Table 2:**
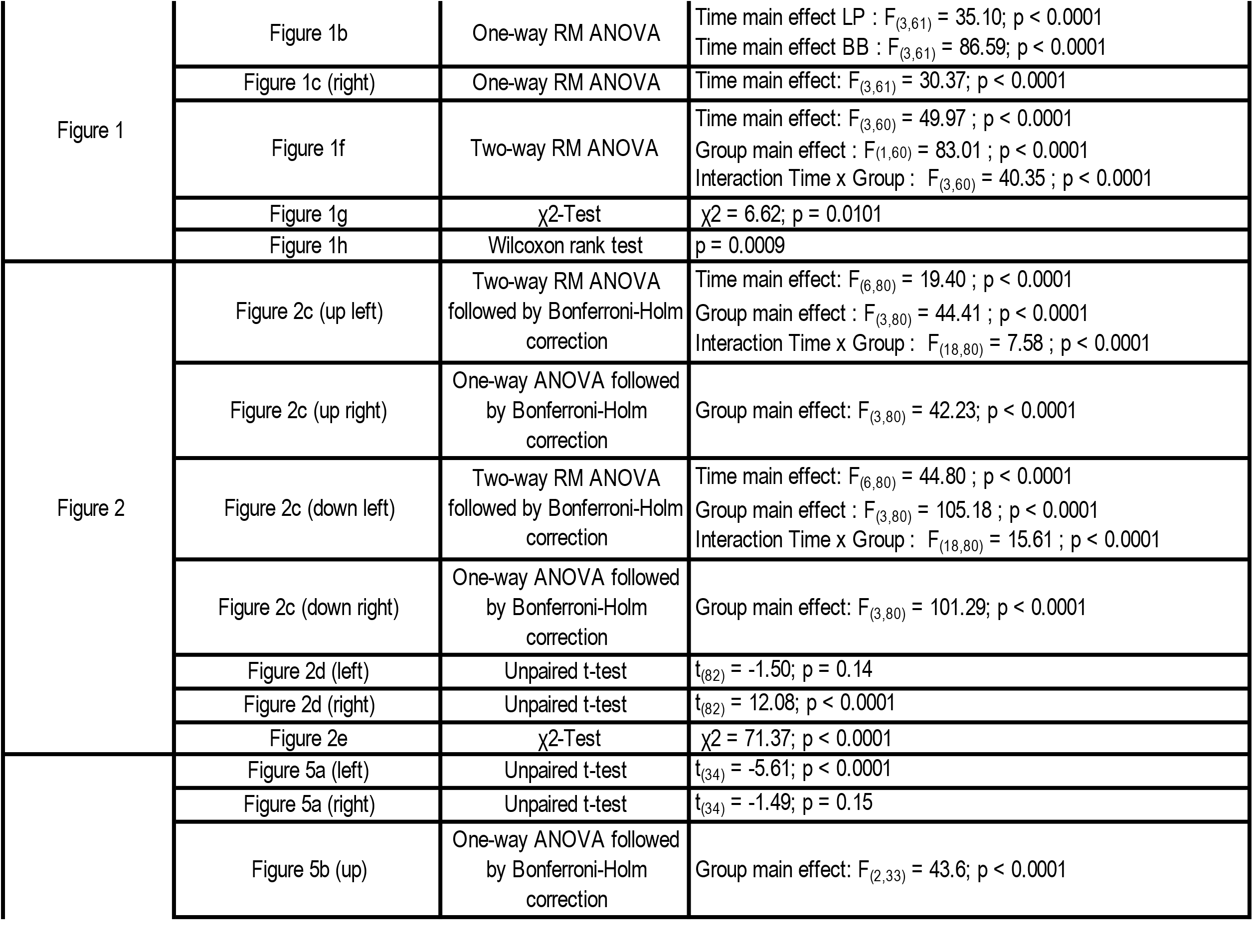
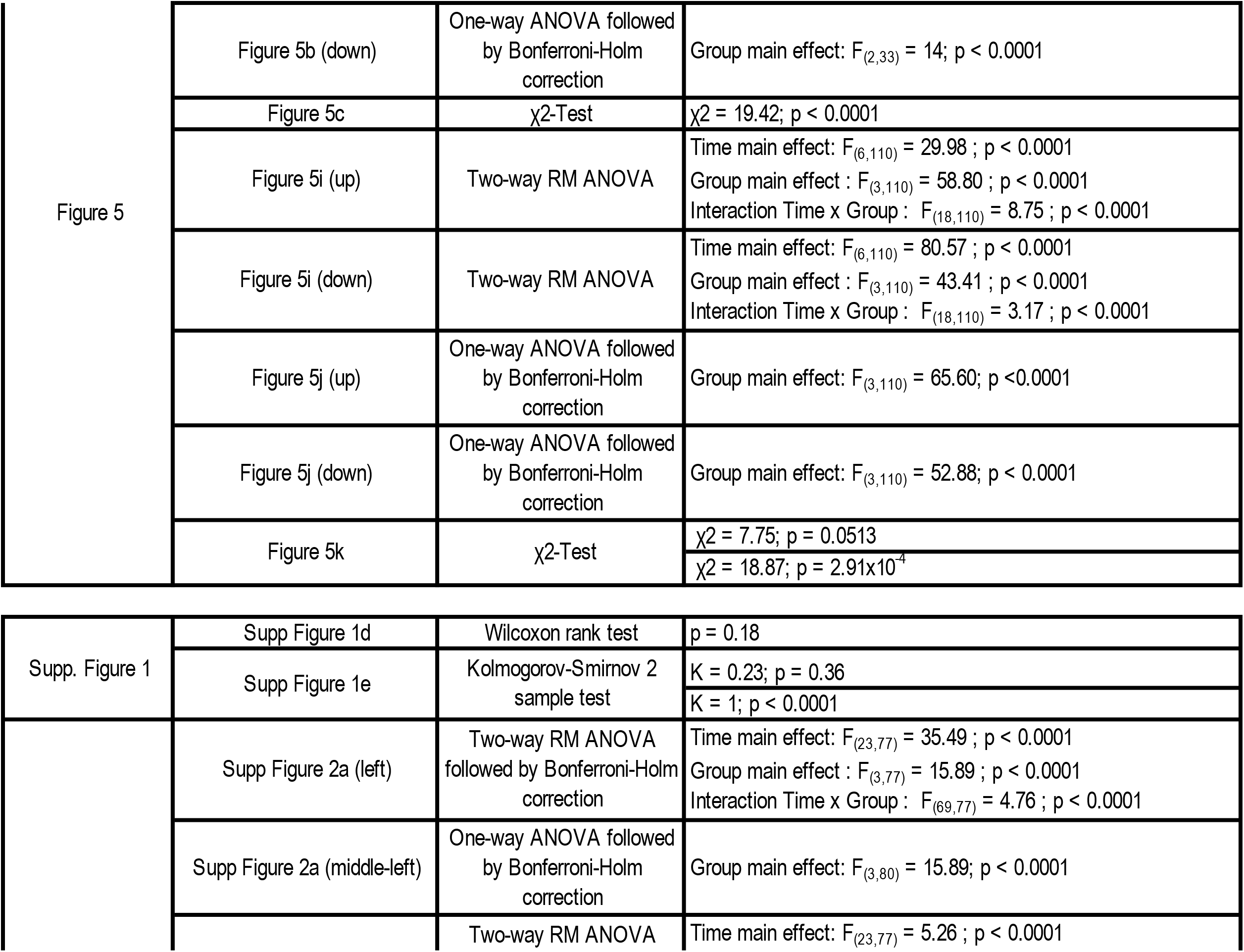
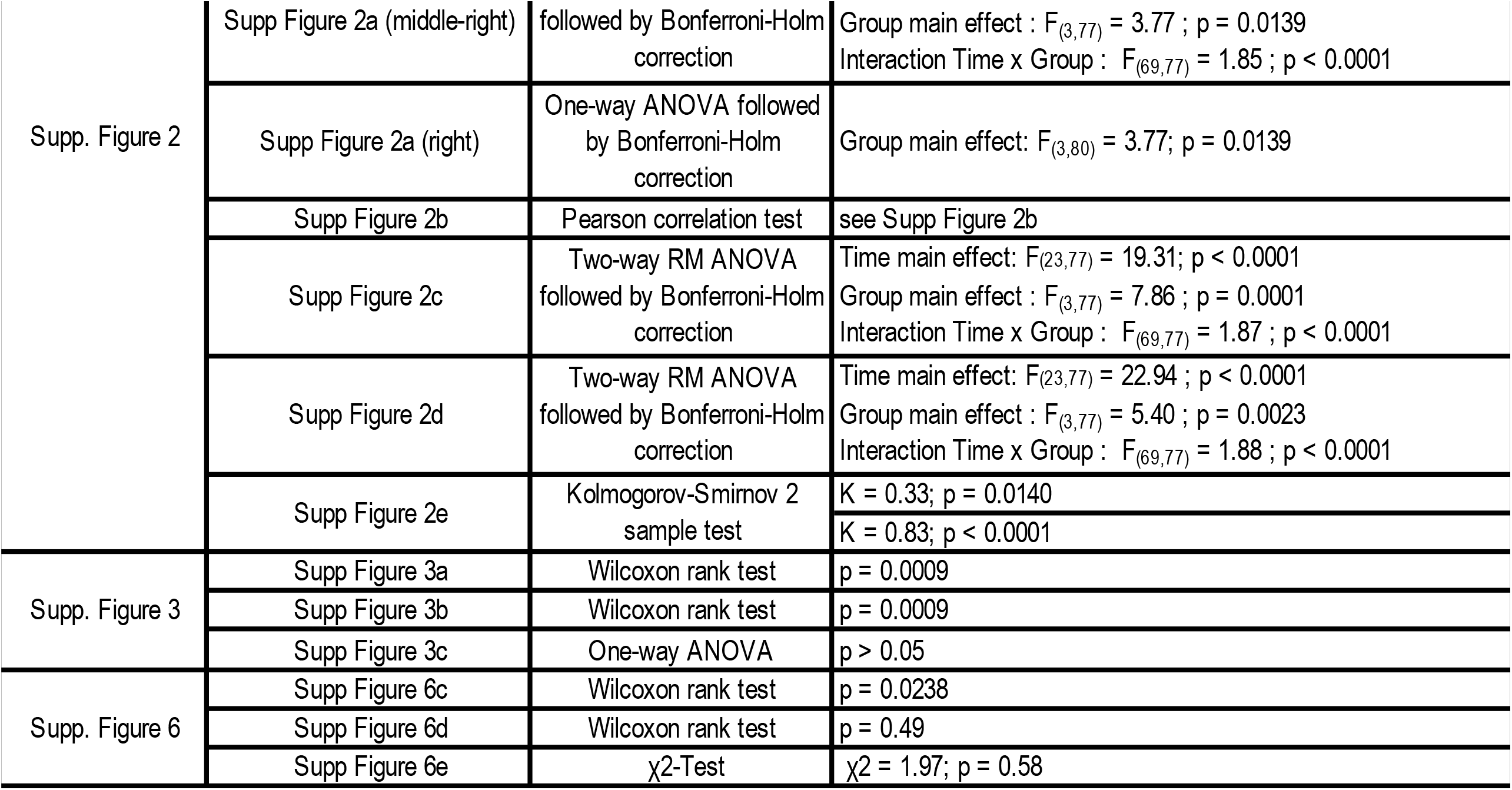
Figures statistical tests.

### Statistics

A priori power analyses were not conducted to predetermine the exact number of animals utilized in the experiments. Randomization was implemented at the point of viral infection or behavioral testing to assign animals to their respective groups. Statistical analyses were performed using MATLAB. The normality of data distributions was assessed using the Shapiro–Wilk test. For data sets failing the normality test, non-parametric statistical tests were employed. Normally distributed data were analyzed using independent or paired t-tests depending on the study design. In cases where data normality was violated, the Mann– Whitney test or the Wilcoxon matched-pairs signed-rank test were implemented. For experiments involving multiple comparisons, repeated-measures (RM) ANOVA was used. RM two-way ANOVA was employed for analyses involving two factors, assuming data normality and applying the Bonferroni–Holm correction post-hoc. To compare proportions and ratios between groups, the chi-square (χ^2^) test was applied. All statistical tests were two-tailed. Data are presented as mean ± s.e.m., with statistical significance established at a p-value threshold of P < 0.05.

## Data and materials availability

A table summary of the experimental data is in the Supplementary Material. Source data are provided with this paper. In addition, some of the data generated in this study will be deposited in the Zenodo database. Full behavioral data generated during and/or analyzed during the current study are available from the corresponding author on reasonable request.

## Code availability

Code for the model and for the data analyses will be made available in the Zenodo database.

## Acknowledgements

We are grateful for support from Otilia de Oliveira, Noemie Karakaplan-Dherbe and Emilie Tubeuf at ESPCI animal facilities. Clément Solié would like to thank “La Fondation des Treilles” for supporting this work.

## Author Contributions

Conceptualization: CS and PF

Experimental Methodology: CS AN RJ FdC CV AM SD ND

Investigation: CS AN BM CB LR TleB SlF AG JAV

Experimental data Analysis: CS BM FM RJ PF

Numerical modelling: BD JN BG YL PF

Reduced model: BD

Funding acquisition: PF BD CS

Writing – original draft: CS BD PF

Writing – review & editing: CS AN YL CV JPH AM JN

## Supplementary Figure legends

**Supplementary Figure 1:**
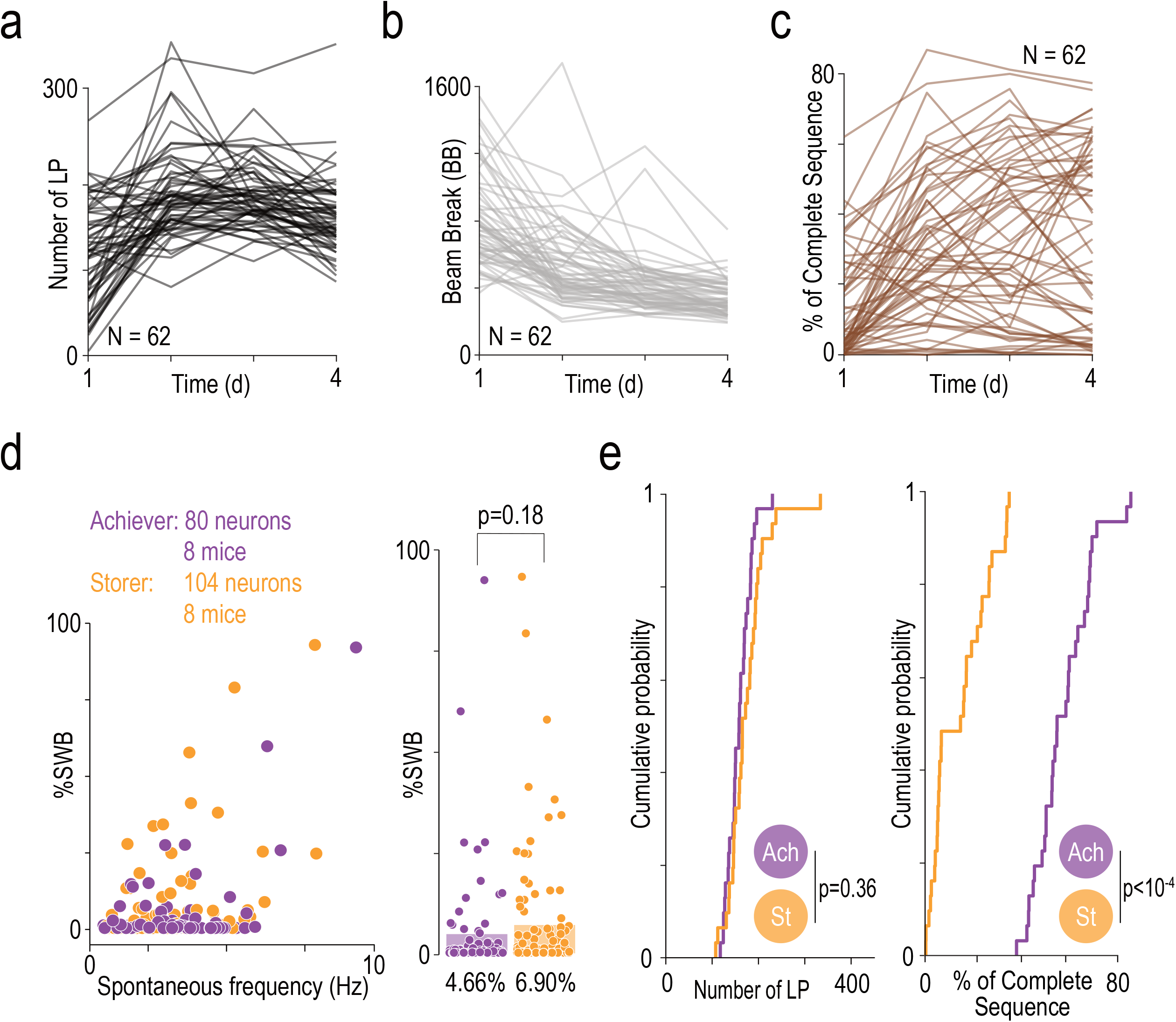
**(a)** Number of lever-presses (LPs) through the different days in lone condition for each animal. **(b)** Time course of the number of beam breaks (BB) for each animal. **(c)** Time course of the percentage of complete sequences for each animal in lone condition. **(d)** In-vivo electrophysiology in anesthetized mice. (Left) Spontaneous frequency in hertz (Hz) in function of the percentage of spikes within burst (%SWB) for each neuron recorded, depending on the profiles of the mice. (Right) Average of the %SWB between *Achievers* and *Storers* male mice. **(e)** Cumulative probability of the number of LPs (left) and of the percentage of complete sequences (right) between *Achievers* and *Storers* mice for the last two days of experiment.

**Supplementary Figure 2:**
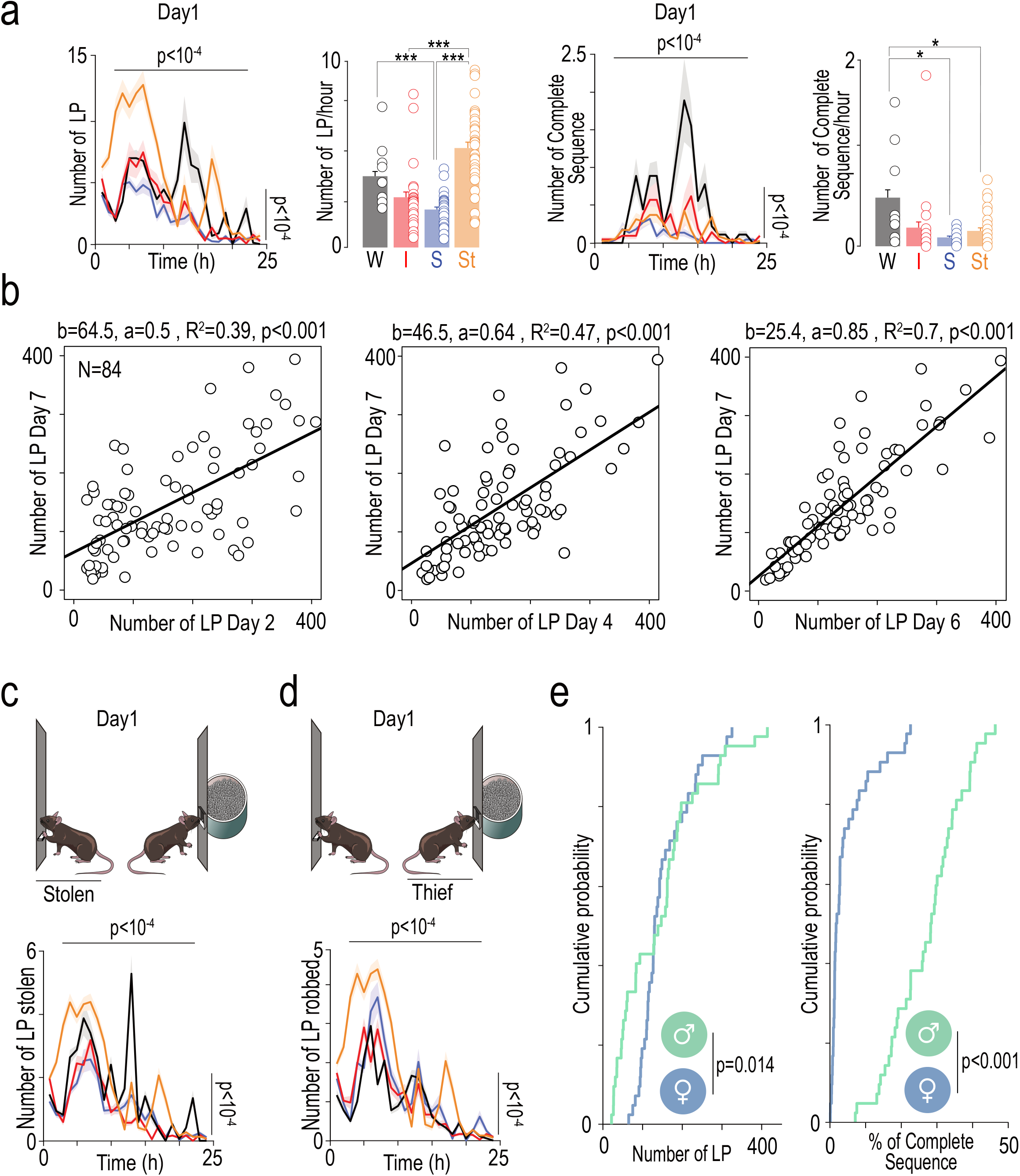
**(a)** (Left) Time course of the number of LPs during the first day for the different profiles of mice in the social condition. (Middle-Light) Average number of LPs per hours the first day between the different profiles. (Middle-Right / Right) Same analysis than described previously for the percentage of complete sequences. Seq. **(b)** Correlations analysis for the number of LPs between the last day and day2 (left), day4 (middle) and day6 (right), for each mouse. **(c)** Number of food pellet stolen to a mouse (Stolen) after an LP during the first day depending on the profile. **(d)** Number of food pellets stolen by a mouse (Thief) after an LP during the first day depending on the profile. **(e)** Cumulative probability of the number of LPs (left) and in the percentage of complete sequences (right) between male and female mice in social condition.

**Supplementary Figure 3:**
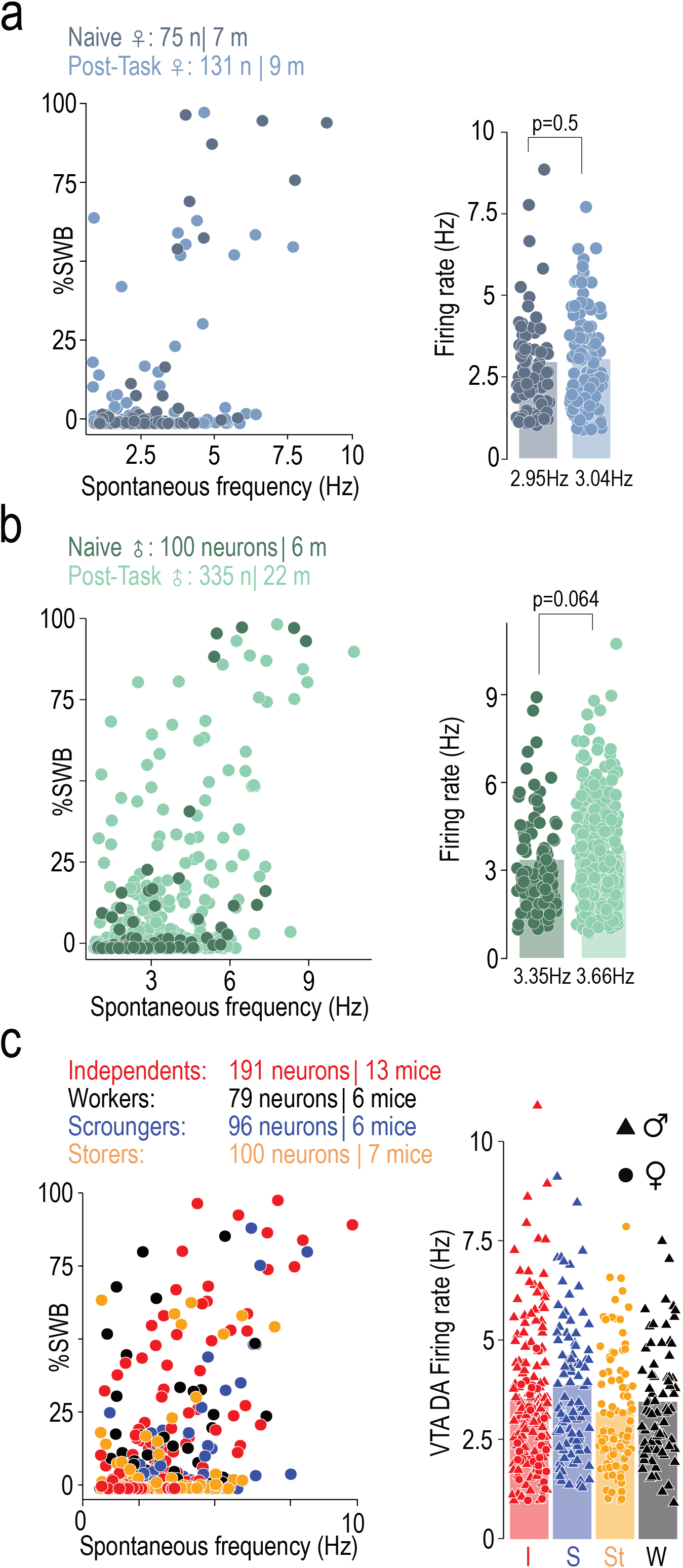
**(a)** In-vivo electrophysiology in anesthetized mice. (Left) Spontaneous frequency in hertz (Hz) in function of the percentage of spikes within burst (%SWB) for each neuron recorded in female mice in naïve condition and after the social condition. (Right) Average of the VTA DA firing rate between naive and after the task in female mice. **(b)** Same as **(a)** for male mice. **(c)** In-vivo electrophysiology in anesthetized mice. (Left) Spontaneous frequency in function of %SWB for each neuron recorded depending on the behavioral profiles after the social condition. (Right) Average of the VTA DA firing rate between the different profiles after the task.

**Supplementary Figure 4:**
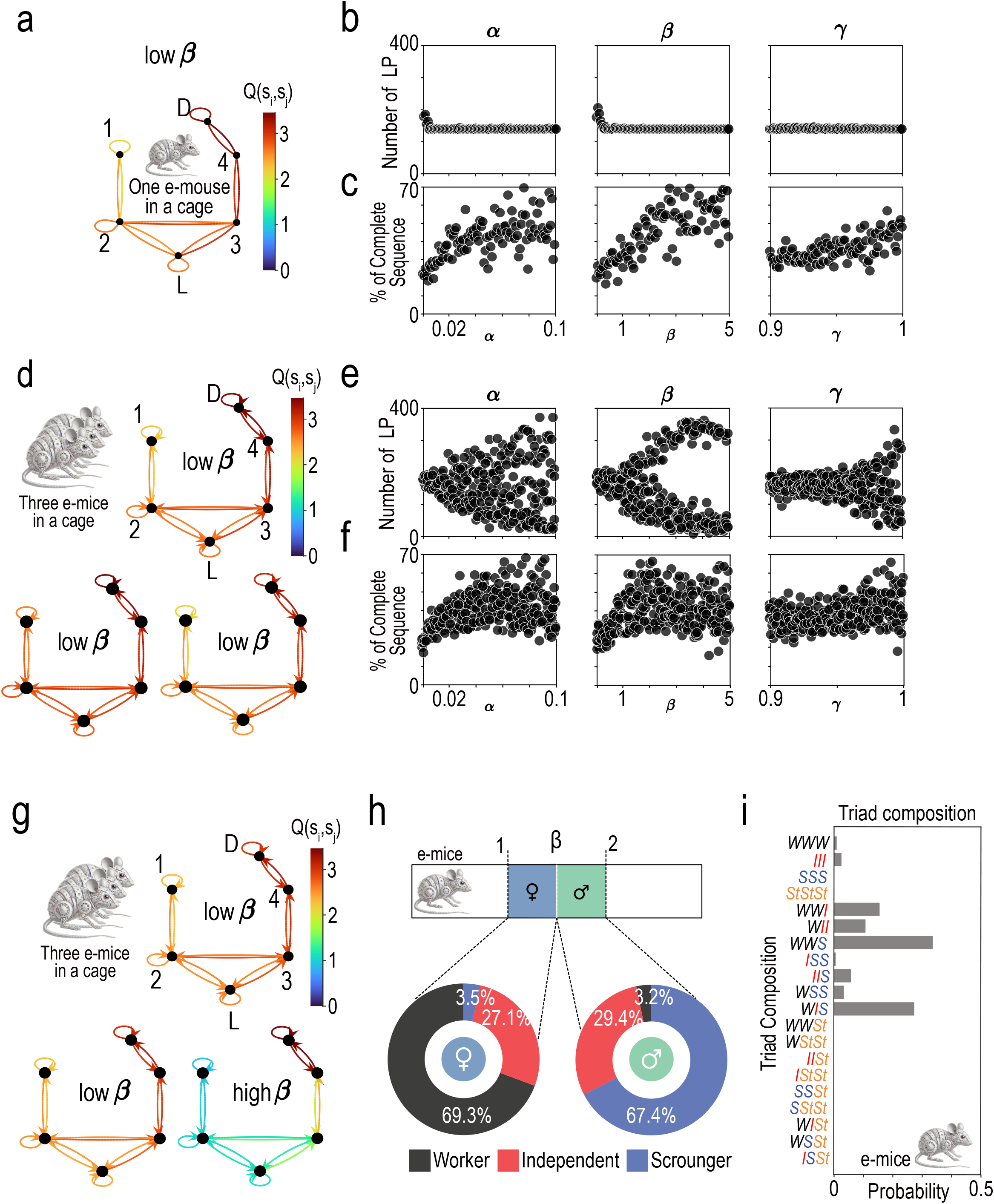
**(a)** Q-learning simulation of one e-mouse with « low » β (β=0.5; compare to Fig. 3a, where β is high). **(b-c)** Dependence of LP count **(b)** and in the percentage of complete sequences on the learning rate (α), inverse temperature (β) and temporal discount factor (<u>γ</u>) in the full model and lone condition. In each panel, the considered parameter is varied, with other parameters set at their standard value (methods). **(d)** Same as **(a)**, with three e-mice with low β (β=0.5; compare to Fig. 3d, where β is high). **(e-f)** Dependence of LP count **(e)** and in the percentage of complete sequences **(f)** on parameters, as in **(b-c)**, but in the social condition, with three e-mice having the same parameter values. **(g)** Same as **(d)**, with two e-mice with a low β and one e-mouse with a high β (compare to **(d)**, with three low β e-mice and Fig. 3d, with three high β e-mice). **(h-i)** Mean proportions of the three profiles **(h)** and of mean probabilities of box types **(i)** of Fig. 3h.

**Supplementary Figure 5:**
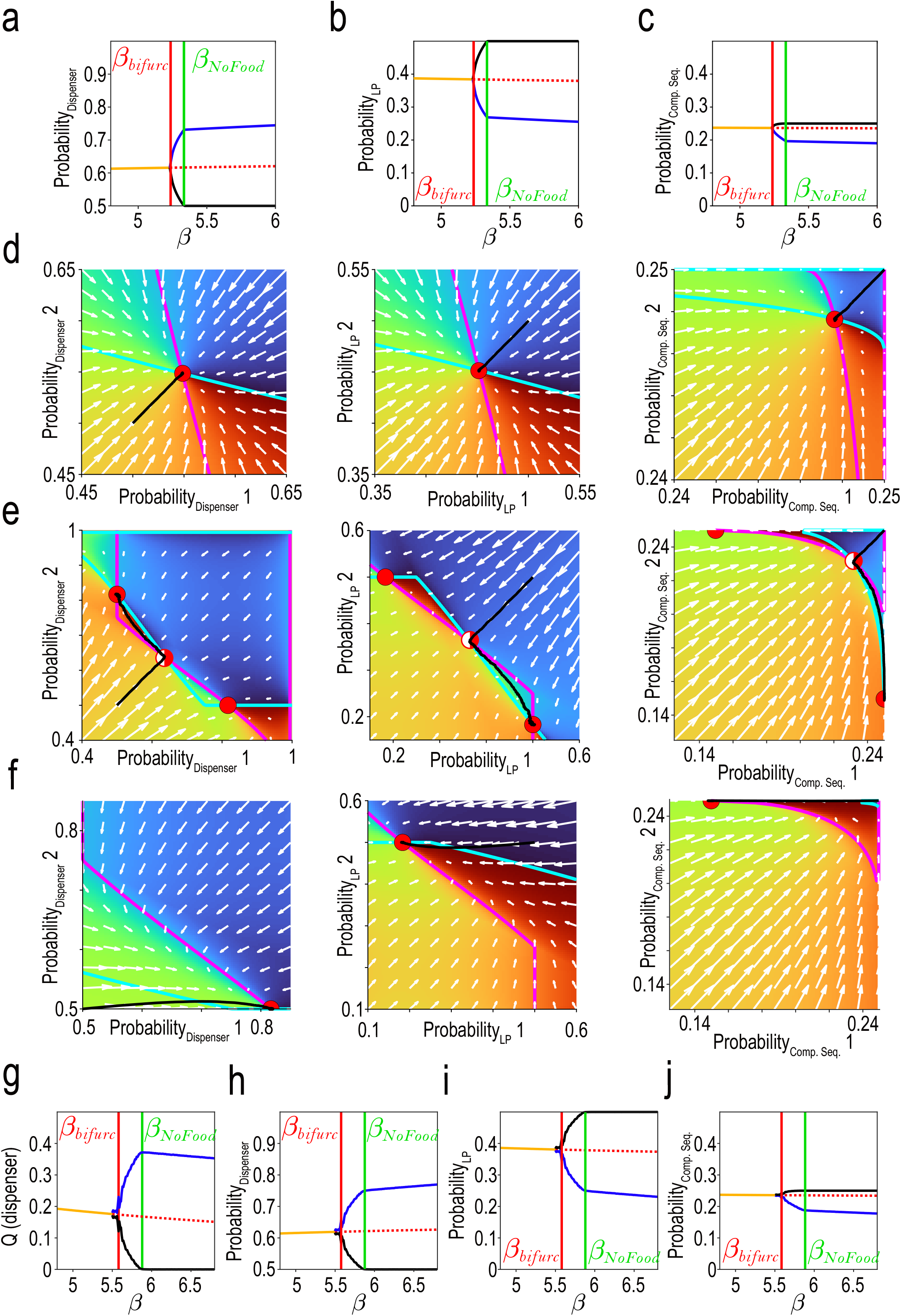
**(a-c)** Bifurcation diagrams corresponding to Fig. 4a, as a function of the residence probability at the dispenser **(a)**, lever press **(b)** and complete sequence **(c)** probabilities in the reduced model. **(d-f)** Model dynamics corresponding to Fig. 4b-c in the residence probability at Dispenser (left), lever press (center) and complete sequence (right) probabilities planes, for females **(d)**, males **(e)** and in the mixed box condition. **(g-j)** Same as **(a-c)** for the reduced model when no linearization of the softmax function is done. The bifurcation diagrams are qualitatively and quantitatively similar to those of the reduced model with linearization (compare to Fig. 4a, right and **(a-c)**).

**Supplementary Figure 6:**
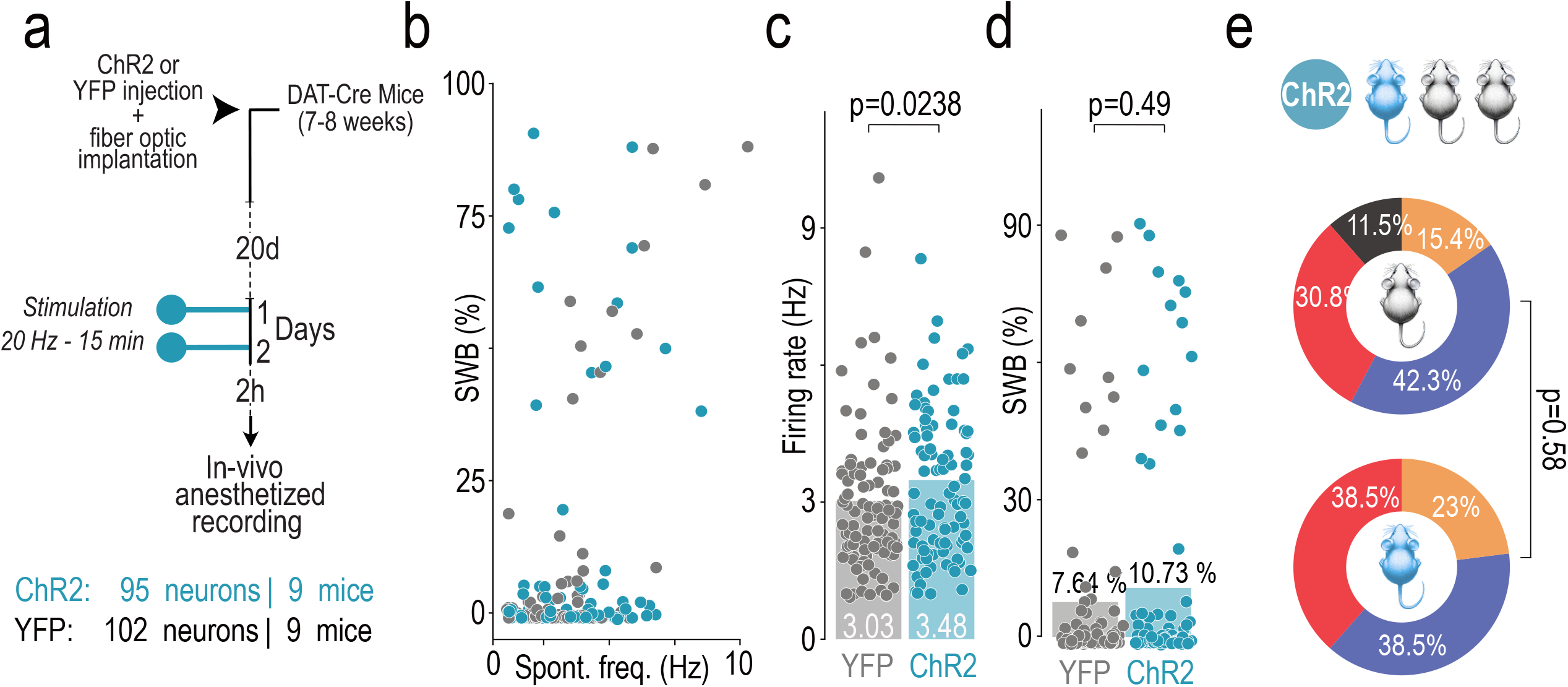
**(a)** Time course of YFP or ChR2 transfection in the VTA DA neurons of DAT-Cre mice and optic fiber implantation, followed by stimulation protocol 24 hours and 2 hours before in-vivo anesthetized electrophysiology. **(b)** In-vivo electrophysiology in anesthetized mice. Spontaneous frequency in hertz (Hz) in function of the percentage of spikes within burst (%SWB) for each neuron recorded in YFP and ChR2 mice after the stimulation protocol. **(c)** Average of the VTA DA firing rate between YFP and ChR2 mice after the stimulation protocol. **(d)** Average of the VTA DA %SWB between YFP and ChR2 mice after the stimulation protocol. **(e)** Comparison of the repartition of the different behavioral profiles between the group of mice expressing YFP and the group of mice expressing ChR2, in triads containing one ChR2 and two YFP mice.

## Modeling (Supplementary Methods)

### Building a behavioral model of e-mice lone and social behaviors

The environment of experiments was modeled as 6 states (Fig. 3 and Supp 4; rooms 1-4, and lever and dispenser positions). The number and sex of e-mice present in the environment was varied, with either 1) one (male or female) e-mouse in the lone experiment, 2) three males or females e-mice in social experiments, or 3) 1 male and 2 females in the mixed-box experiment. In the model, we considered e-mice satiety, to account for the convergence of pellets’ consumption, *n*_*Eat*_, toward a maximum, *n*_*Sat*_. Motivation was considered to decrease with satiety:

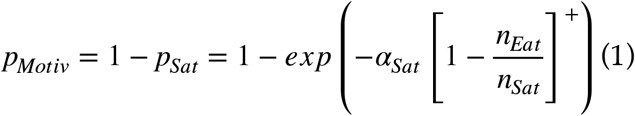

and scaled both action probabilities and the learning rate (see below). Moreover, e-mice were considered to encounter both pressing and eating fatigue in the model, to get paces consistent with those observed in real mice (without fatigues, paces were unrealistically high in the model). Both fatigue processes were described as action probabilities decreasing upon each individual action and restoring back to their maximal value with first-order dynamics :

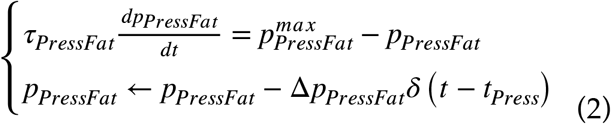

and

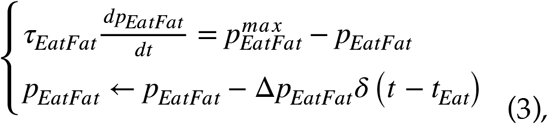

with *δ* being the Dirac delta function. Both fatigue processes and satiety concurred to decrease action probabilities in the model. When at the lever, e-mice pressed the lever with probability

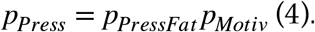

The lever could not be pressed for 3 time steps after each press (i.e., to mimic the 5s lever unavailability in experiments). When at the dispenser, e-mice ate with probability:

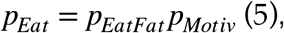

when pellets were present. State transition occurred at each time step, with softmax probabilities based on *Q* values of all accessible *k* states from the current state *s*_*i*_:

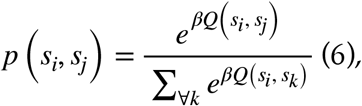

with *β* the inverse temperature parameter favoring either exploration (i.e., random choices; lower *β*s) or exploitation of previous learning (higher *β*s). *Q* learning occurred upon each transition from a departure state *s*_*i*_ to an accessible arriving state *s*_*j*_, as

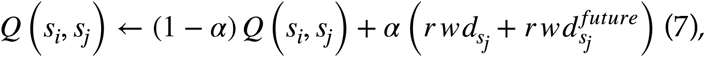

with *α* the learning rate, 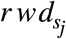 the reward obtained at the arriving state *s*_*j*_ and 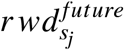 the maximal expected reward obtainable in states accessible from the arriving state *s*_*j*_:

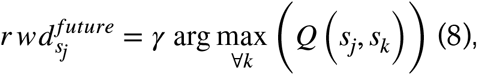

*γ* being a temporal discount factor. The learning rate scaled with motivation, which was considered to decrease with satiety,

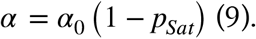

As a consequence, learning stopped when e-mice were satiated. The *Q* matrix was initialized with value 1 at all elements corresponding to possible transitions (and 0 elsewhere). Consistent with observed trajectories in real mice, stops were forbidden at rooms 3 and 4, i.e., on trajectories back and forth between lever and dispenser states.

### Observables

The number of lever presses was computed – for each e-mice – from the beginning of the simulation up to its satiety. The percentage of complete sequences was computed as the number of direct sequences from lever to dispenser (within 3 times steps, i.e. 6 seconds, as in experiments), normalized by the number of lever presses:

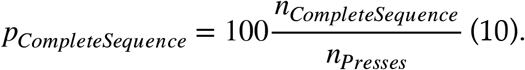

These two observables are depicted in **Fig. 3a, e** and **h** and were used to categorize e-mice and e-mice triads, as done in experiments (**Fig. 3c, e, g, h, i**). In each experimental condition (alone, social, mixed), a hierarchical clustering of e-mice was achieved with Matlab linkage method, based on the number of lever presses and the percentage of complete sequences, after removal of outliers on both variables (5% bilaterally). This procedure was essential as outliers commonly led to unbalanced clustering. This occurred in cases of greatest differences in random realizations of *β* values between e-mice, leading to extreme behaviors (i.e., very « worker » and « scourger » behaviors for very low and high *β*s, respectively), which induced irrelevant clusters including one or a few outliers. Also, both variables were normalized before clustering:

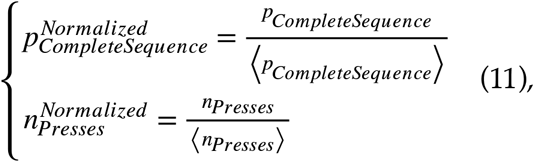

so that the clustering was influenced equally by both variables.

### Reduced model of social interactions

We built a reduced theoretical model (hereafter termed the *reduced model*) to assess, within a mathematically tractable framework, the causal mechanisms whereby specialized behaviors emerge under social interactions. To do so, we derived the reduced model from the full one, based on a continuous time version of *Q* dynamics (i.e., ordinary differential equations, ODEs). In these EDOs, learning and behavioral dynamics operated at a slower time scale – compared to that of individual choices in the full model – so that actions (i.e., state transitions, pressings, eatings) were described probabilistically in the reduced model. In this framework, we performed qualitative analysis of ODEs to determine the number and stability of fixed points of learning and behavioral (state) variables as a function of parameter (with as focus on *β*, which is essential in setting social interactions in the full model). Moreover, to reduce dimensionality for better tractability, we considered a simpler setup, where the environment contains only two positions for e-mice (the lever,*L*, and the food dispenser, *D*), only two e-mice (which allowed to assess social interactions) and we did not consider fatigue or satiety. Here, *q* dynamics were derived from equation (7), to account for their convergence to the expectation of present and future rewards (as for the dynamic of their discrete time equivalent *Q* values in the full model). Hence,

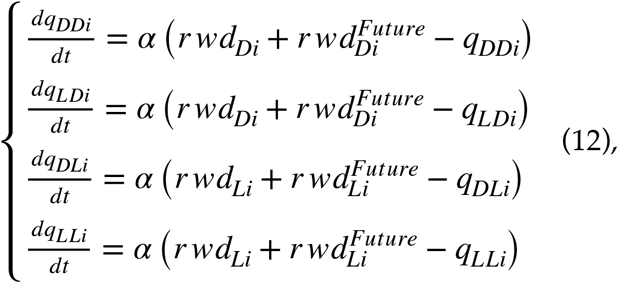

with *q*_*XYi*_ being the *q* value of *X* →*Y* transition, and *r wd*_*Xi*_ and 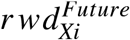 the present and future expected reward at position *X* in e-mouse *i*. Note that, despite all the simplifications made, the dynamical system under consideration is still 8-dimensional. Fortunately, it is straightforward to note that dynamics of *q*_*DDi*_ and *q*_*LDi*_ are identical (as that of *q*_*LDi*_ and *q*_*LLi*_), so that, starting from identical null initial conditions

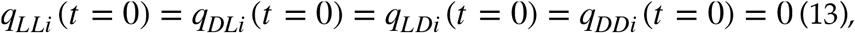

one has

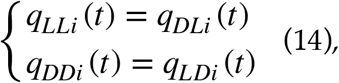

so that only one *q* value at each position is sufficient to describe dynamics:

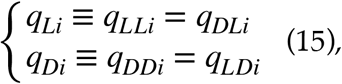

reducing the theory to a 4-dimensional problem. For each e-mouse, one could express *q* values evolved according to :

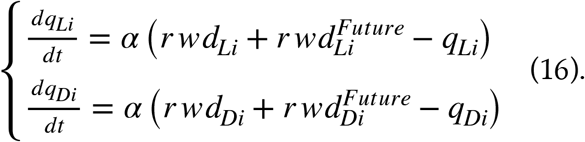

The mean expectation of current reward at *L* was null (*r wd*_*Li*_ = 0), while that at *D* scaled with the probability of being at the dispenser, the probability of eating, the probability of available food at the dispenser, and the reward value of eating, which writes, for e-mouse #1, e.g.,

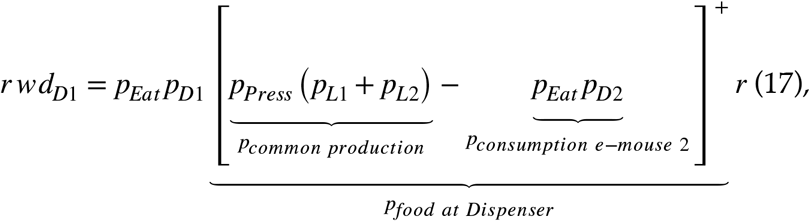

with *p*_*Xi*_ the residence probability of being at position *X* for mouse, *p*_*Eat*_ and *p*_*Press*_ the probabilities of eating and pressing, and *r* the unit reward per eating. The positive part function,[]^+^, is necessary to account for a positive probability of available food at the dispenser. Note also that because *p*_*Di*_ > 0.5 and *p*_*Li*_ = 1 − *p*_*Di*_ < 0.5 (as there is a reward at *D* and not at *L*), the probability of available food at the dispenser is inferior to 1. Note that equation (17) can be written for mouse #2’s by swapping indices #1 and #2, as can be done for other equations in the following. Whenever possible, we keep the generic index *i* for e-mice. By else, at both positions, the expectation of future rewards expressed as

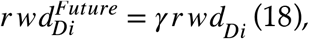

as getting a reward at *D* is the maximal expected possible reward at the next step from both positions *L* and *D*. Therefore, *q* derivatives can be re-expressed as

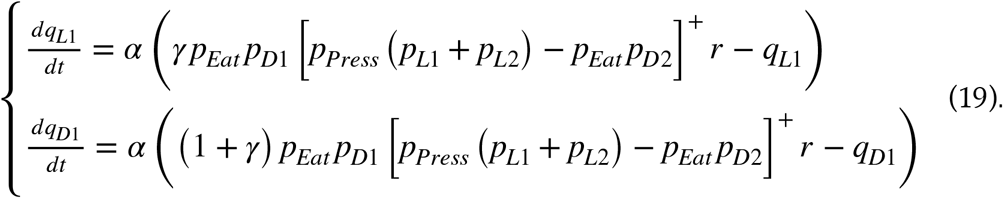

In the following, we considered, for the sake of simplicity that *p*_*Press*_ = *p*_*Eat*_ = 1. Also, because *p*_*Li*_ = 1 − *p*_*Di*_ for both e-mice *i, q* derivatives simply expressed as

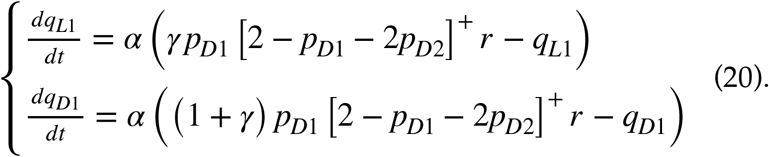

Note that the first equation is actually the second one scaled by a factor 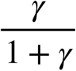. As both *q*_*D*1_ and *q*_*L*1_ both start from null initial conditions (equation (13)), their dynamics are homothetic, so that at any point in time

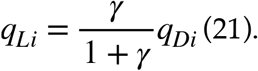

In the following, we proceed with *q*_*D*1_ equation, which, together with that for *q*_*D*2_, describes the system with two-dimensional dynamics:

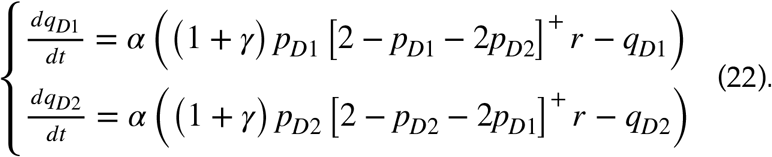

In such a setup, with two states, residence probabilities of e-mouse *i*, i.e., *p*_*Li*_ and *p*_*Di*_ can be directly related to transition probabilities. Indeed, assuming first-order state transitions, *p*_*Di*_ evolves as:

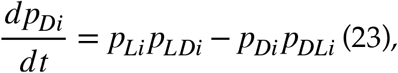

which, after substituting *p*_*Li*_ by 1 − *p*_*Di*_ leads to

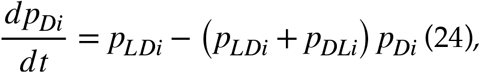

with a unique fixed point

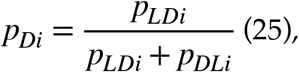

so that replacing *p*_*LDi*_ and *p*_*DLi*_ by their softmax expression, leads to

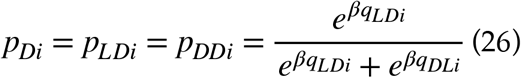

and

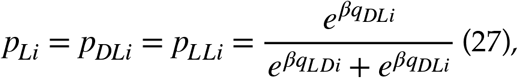

which, thanks to equation (15), can simply be written

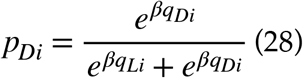

and

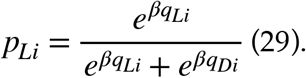

Combining equations (21) and (28) allowed to express the residence probability as a function of the *q* value at dispenser:

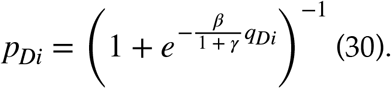

Then, combining equations (22) and (30) yields

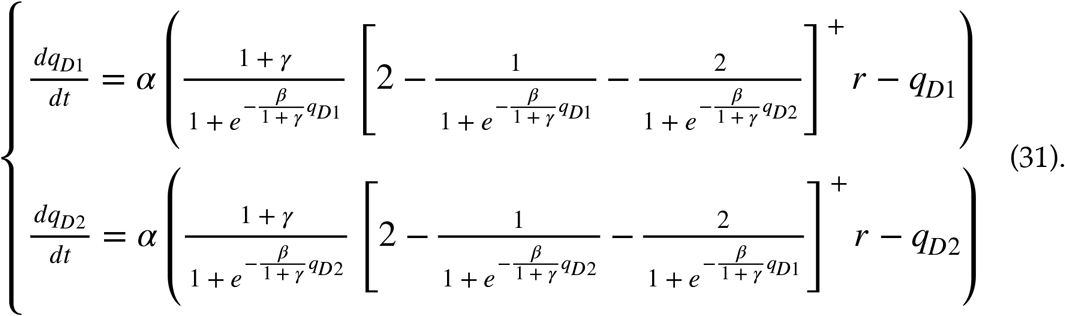

Note that this system describes the social interaction between two e-mice having similar *β* values (I.e., two males or femelles). This two-dimensional EDO system describes the time evolution of learning and behavior variables (as residence, lever press and direct sequence probabilities can be derived from *q* values, see below). Unfortunately, this system (even studied by parts, depending on whether probabilities of food at dispenser ate positive or null) has no analytical solution. We therefore assessed a version of the EDOs with linearized approximations of residence probabilities. This linearized version could be solved analytically, which allowed to assess the number and stability of fixed points as a function of *β*. To do so, we used a linearized approximation of residence probabilities at position *X* in e-mouse *i*, i.e.

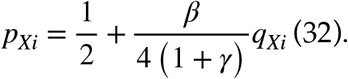

Such approximation was typically of the order of one to a few percents. Combined with equation (22), it allowed to provide the tractable system we analyzed to generate **Figs. 4 et Supp 5** and which, after a few lines of algebraic manipulation, expresses as:

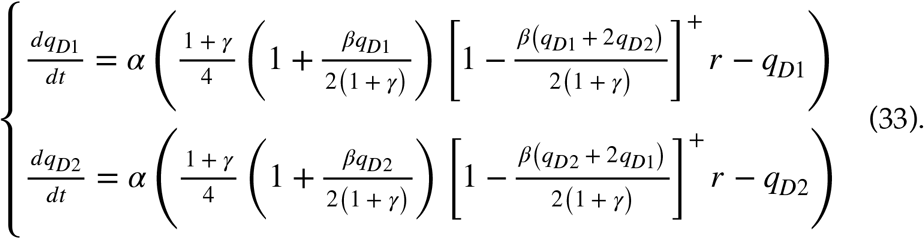

In the reduced model, the propensity of e-mouse *i* to press the lever was expressed as a probability, which scaled both with the probability to be at the lever position and the probability to press, once at that position:

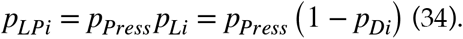

The propensity to achieve direct sequences was also expressed as a probability, scaling both with the be at the lever position, to press the lever and then move toward the dispenser position (which is equal to the residence probability at dispenser (equation (26)):

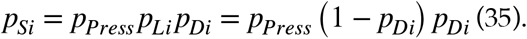

### Simulation of the reduced model

The reduced model was simulated using a noisy version of system (19):

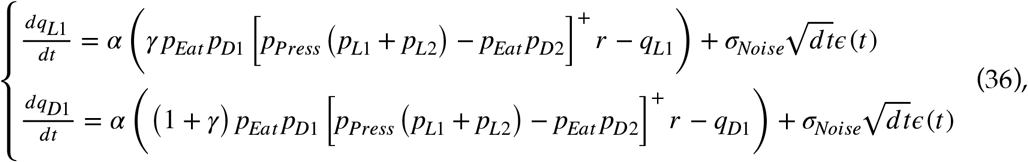

where *σ*_*Noise*_ is the noise standard deviation and *ϵ* (*t*) is a Gaussian stochastic process with zero mean and unit standard deviation, similar equations for e-mouse #2 and residence probabilities computed according to equation (30). Simulations were used to check the validity of the qualitative analyse achieved to produce bifurcation diagrams and illustrate system’s dynamics in phase portraits (see below).

### Qualitative analysis of the reduced model

The system (33) admitted one or several fixed points 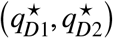, which number and nature depended on parameters. We focused on the dependence of 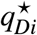 steady-states of individual e-mice on *β*, i.e. 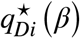, denoted *branches* in the following (see figures Xxxx). Indeed, as in the full model, the value of *β* separated distinct regimes of social interaction between e-mice. The main results of the qualitative analysis of the system are recapitulated in Table 1.

We found that for *β* values below a critical value *β*_*Bifurc*_ (see below, for the analytical calculation of critical values), only one fixed point, 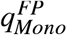, existed, with both e-mice admitting a similar steady-state:

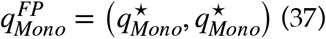

with

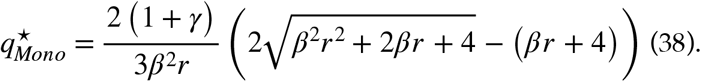

Its stability was set by

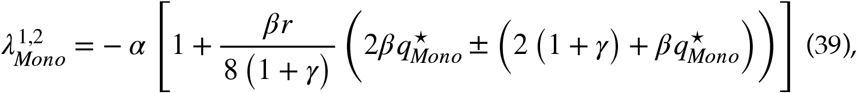

with both Eigen values negative, so that 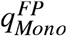 was a stable node.

Above 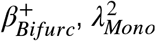, was positive, so that 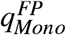 became a saddle node. Also, two other symmetric fixed points appeared, 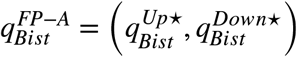 and 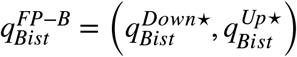, with

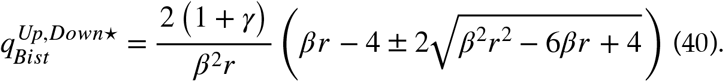

The fixed point 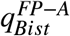 corresponded to the case in which e-mouse #1 displayed a higher *q* value at dispenser 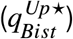, and, as a consequence, a higher residence probability 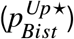 at dispenser, and lower probabilities of lever presses 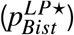 and direct sequences 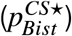, compared to e-mouse #2 (see below). In other words, 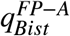 was a fixed point at which e-mouse #1 behaved as a scrounger and e-mouse #2 as a worker, while behaviors were inverted at 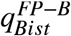. In this context, the 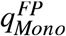 saddle node branch could be interpreted in this reduced model as corresponding to independent e-mice found in social male boxes in experiments and the full model (starting from (0,0) with small levels of noise yields animals to this saddle branche, see below). The stability of both fixed points was set by

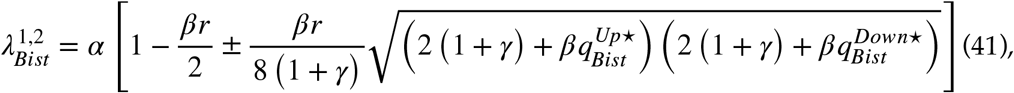

with both Eigen values negative, so that both 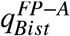 and 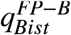 were stable nodes.

For *β* values above a second critical value *βNoFood* (with *β*_*NoFood*_ > *β*_*Bifurc*_, see below), the probability of available food at the dispenser became null for the worker e-mouse (i.e., the argument within the positive part function, []^+^, becomes negative for e-mouse #1 or 2 in equation (33)). The system thus writes, if the worker is e-mouse #2,

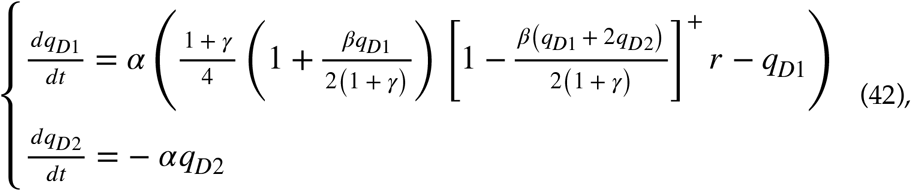

A single fixed point 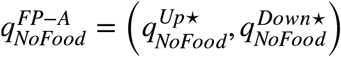 existed

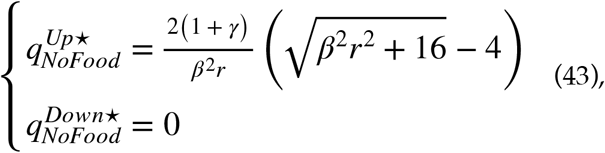

which stability was determined by

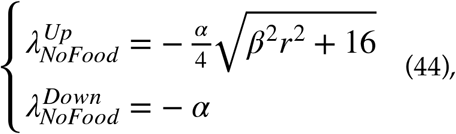

with both Eigen values negative, so that 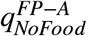 was a stable node. Obviously, a symmetrical stable node, 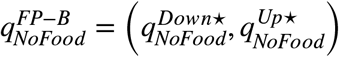, co-existed with 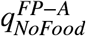, when the worker was e-mouse #1.

Both *β*_*Bifurc*_ and *β* _*NoFood*_ critical values could be found analytically. At the bifurcation point where 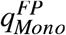 becomes a saddle node, the two symmetric fixed points 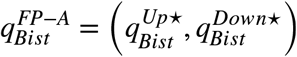 and 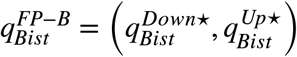 switch from complex (*β* < *β*_*Bifurc*_) to real (*β* ⩾ *β*_*Bifurc*_) values. Thus, this point was set by *β* by a simple criterion, i.e., canceling of the term under the square root function in equation (40):

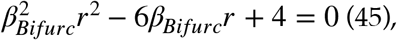

which yields

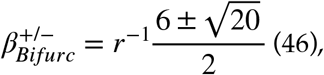

with 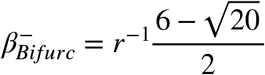 separating branches of real negative 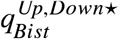 values from the branch of complex values between 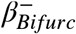 and 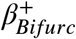. The branch of real negative 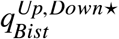 values below 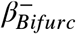 corresponded to impossible solutions, as *q* values are positive (with a null initial condition (equation (13)) and positive reward expectations). Finally,

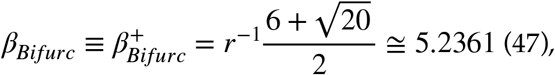

was the critical point (assuming *r* = 1; see parameter values, below) at which the system switched from one to three fixed points (two stable ones separated by an unstable one), at a supercritical pitchfork bifurcation (« trifurcation »; Strogatz, 1994). At this point, where 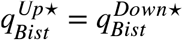, i.e., at which 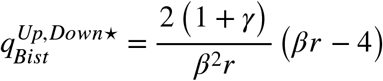, one could easily check, using equation (39), that 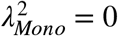 i.e. 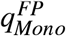 looses its stability along the second Eigen direction (i.e. switching from a stable to a saddle node).

The other critical value, *β*_*NoFood*_, could be found by considering that 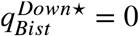 at that point, i.e.,

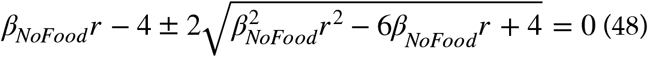

which yields

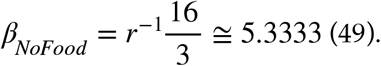

Bifurcation diagrams were built (**Figs. 4 et Supp 5**) by representing 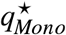 stable and saddle branches (below and above *β*_*Bifurc*_, respectively, 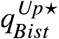 and 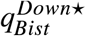 branches for *β*_*Bifurc*_ ⩽ *β* < *β*_*NoFood*_, and 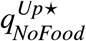 and 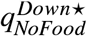 branches for *β* ⩾ *β*_*NoFood*_.

We verified that bifurcation diagrams obtained using linearized approximation of residence probabilities (equation (32)) were indeed qualitatively correct and quantitatively consistent with the original reduced model, using numerical simulations of the reduced model. To do so, the stable and bistable branches were computed by taking steady-state *q*_*Bi*_ values of these simulations, and computing *p*_*Bi*_, *p*_*LPi*_, and *p*_*Si*_ from equations (30), (34) and (35), respectively. The 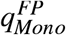 saddle node branch was acquired by doing the same, in the absence of noise (*σ*_*Noise*_ = 0), so that system’s state converged to the saddle node from its null initial condition (i.e., along the first bisector in the *(q*_*B*1_, *q*_*B*2_) plane, due to system’ symmetry) and stayed at it, without further diverging toward 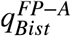 or 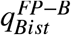 (which occurred in presence of noise), as Eigen directions are stable along the first bisector and unstable along the second bisector (see below).

### Eigen Vectors of the reduced model

Eigen Vectors in the general case expressed as

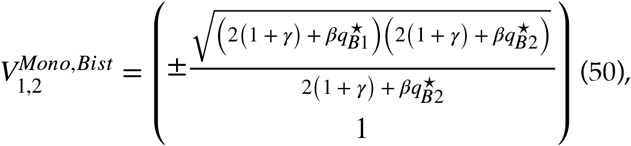

which, in the case of the 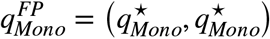 branch, trivially simplified to

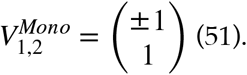

In the case with no food for one e-mouse, Eigen Vectors expressed as

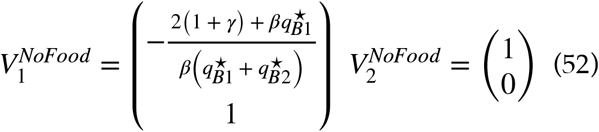

which, in the case of the 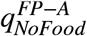, simplified to

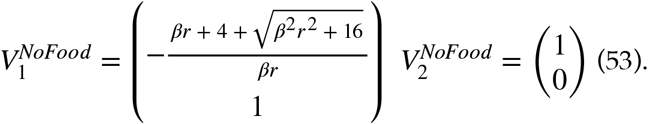

### Phase portraits of the reduced model

In phase portraits (**Figs. 4 et Supp 5**), the *q* vector field was numerically computed from equation (19), with *q*_*Di*_ values set within a fixed range and *q*_*Li*_ computed from equation (21).

The *p* vector field was computed with *p*_*Di*_ values set within a fixed range, *q*_*Di*_ values computed as

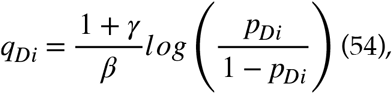

i.e., the reciprocal equation of equation (30), *q*_*Li*_ computed from *q*_*Di*_ with equation (21) and *p* derivatives computed thanks to the chain rule, i.e.,

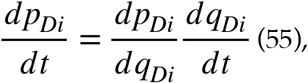

where

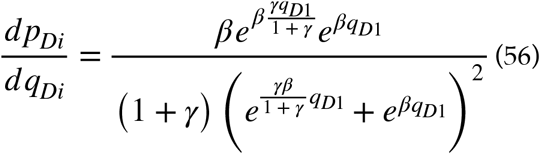

was obtained by differentiating equation (30) and 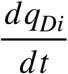 taken from system (19).

The *p*_*LP*_ vector field was computed with *p*_*LPi*_ values set within a fixed range, *p*_*Di*_ values computed as

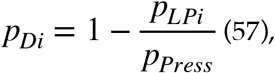

which was derived from equation (34), *q*_*Di*_ values with equation (21),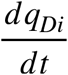 from system (19), 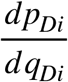 from equation (56) and by applying the chain rule, i.e.,

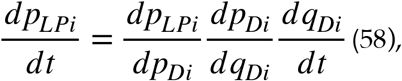

with

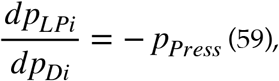

which was obtained by differentiating equation (34).

The *p*_*S*_ vector field was computed as the *p*_*LP*_ vector field, using equations (34), (21), (19) and (56) and applying the chain rule

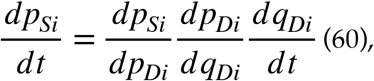

with

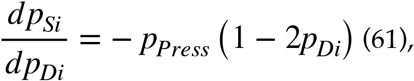

which was obtained by differentiating equation (35).

In phase portraits, vector fields were represented both by a direction field plot and through a color coded map. Nullclines were estimated as zero level contours of the vector fields. Fixed points were taken with bistable Up and Down steady-state values obtained from the bifurcation diagram for *β* = *β*_*Female*_ (social female condition), *β* = *β*_*Male*_ (social male condition). In the mixed box condition, we considered one female and one male and the fixed point was taken as the intersection of both nullclines. Eigen vector were not represented for clarity. An example of simulation of reduced model’s dynamics was also represented in each condition to show dynamics from initial conditions toward the – or one of the – fixed points.

### Numerical procedures

Numerical simulations and qualitative analysis were achieved thanks to Matlab custom scripts that are available online. The qualitative analysis of the reduced model was done using the Symbolic toolbox of Matlab. Numerical simulations of the full and reduced model were performed used Euler forward integration. In the full model, standard parameters, unless mentioned, were *α*_0_ = 0.05, *γ* = 0.99, *α*_*Sat*_ = 20, *n*_*Sat*_ = 150, *τ* _*PressFat*_ = 50*s*, 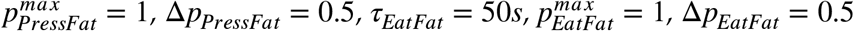, Δ*p*_*EatFat*_ = 0.5, and the time step was Δ*t* = 2*s*. Simulations lasted *t*_*max*_ = 20000*s*. *β* distributions are discussed in the main text. Parameters of the reduced model were as following : *α*_0_ = 0.01, *β* = [1,10], *β*_*Female*_ = 1, *β*_*Male*_ = 10,*γ* = 0.99,*r* = 1,*p*_*Press*_ = 1, *p*_*Eat*_ = 1, *σ*_*Noise*_ = 1*e*^−4^, *dt =* 1*s*. Simulations lasted *t*_*max*_ = 1500*s* for exemple simulations shown in phase portraits and *t*_*max*_ = 100000*s* in simulations with noise, i.e., a much longer time to estimate steady-states in the numerical estimation of phase portraits. The distribution of e-mice behavioral profiles and box type distributions were obtained on 3000 repetitions of 16 boxes of either the lone condition, or single sex or mixed sex conditions, with 16 animals per box (**Fig. 3c, g**, and **Supp. 4 h, i**).

## Extended Data

